# High Connectivity, Local Signatures: Genomic and Otolith Evidence from Reef-Associated Snappers

**DOI:** 10.64898/2026.07.18.735000

**Authors:** Adela Roa-Varón, Nancy G. Prouty, Amanda W.J. Demopoulos, Penny J. McCowen, Laura Thornton, Stacey Harter, Andrew W. David, Santiago Herrera, Andrea M. Quattrini

**Affiliations:** Smithsonian Institution, National Museum of Natural History, Washington, DC, USA; U.S. Geological Survey Pacific Coastal and Marine Science Center, Santa Cruz, CA, USA; U.S. Geological Survey Wetland and Aquatic Research Center, Gainesville, FL, USA; University of Miami Cooperative Institute for Marine and Atmospheric Studies, Miami, FL, USA; NOAA-National Marine Fisheries Service, Panama City, FL, USA; Lehigh University, Bethlehem, PA, USA

## Abstract

Connectivity in reef-associated fishes emerges from processes operating across multiple spatial and temporal scales, yet these processes are rarely evaluated jointly. Here, we integrated genome-wide single-nucleotide polymorphisms (SNPs) data and otolith elemental signatures to examine connectivity in Red Snapper (*Lutjanus campechanus*) and Vermilion Snapper (*Rhomboplites aurorubens*) across 14 reefs and banks from the Flower Garden Banks National Marine Sanctuary and neighboring banks to the east. We tested whether genomic variation revealed population structure across geographic and depth gradients (shallow, upper mesophotic, and lower mesophotic), whether otolith chemistry captured fine-scale site-specific patterns of environmental exposure and habitat use not resolved by genetic markers alone, and how these complementary signals related to spatial variation in age structure and growth dynamics. Genome-wide SNP data revealed little detectable spatial genetic structure across the sampled banks in either species. In contrast, otolith microchemistry revealed pronounced site-associated environmental structuring, indicating that individuals experience localized habitat conditions within a regionally connected system. Red Snapper exhibited otolith signatures consistent with reef-associated environments, whereas Vermilion Snapper showed more variable elemental signatures, indicating exposure to a broader or more heterogeneous range of habitats. Spatial variation in age structure and growth provided additional demographic context, indicating that local differences among banks can emerge despite weak genomic differentiation. Together, these results show that fine-scale ecological and demographic differentiation can exist within a genetically connected system and that otolith chemistry provides critical information on post-settlement processes at scales not captured by genetic markers alone. By linking regional genomic connectivity with otolith-derived habitat signatures and demographic variation, this integrative framework provides a more comprehensive basis for understanding resilience and informing fisheries management in shelf-edge reef systems.

## 1. Introduction

Connectivity is a fundamental ecological process shaping population dynamics, gene flow, and adaptive potential in marine ecosystems. Patterns of connectivity influence species persistence, recovery following disturbance, and responses to environmental change across ecological and evolutionary timescales (Magris et al., 2014; Sahyoun et al., 2016; Harrison et al., 2020). In natural fish populations, connectivity plays a central role in larval dispersal, recruitment success, ontogenetic habitat shifts, and population viability, particularly in heterogeneous seascapes subject to environmental variability and increasing anthropogenic pressures (Fodrie et al., 2020; Vaz et al., 2023). Recent integrative approaches combining genomic, biophysical, geochemical, and ecological data have significantly increased the capacity to characterize these processes, providing a more robust foundation for fisheries management and conservation planning (Cowen & Sponaugle, 2009; Olds et al., 2016; Brown et al., 2016; Bernatchez et al., 2017; Krueck et al., 2022).

Given their ecological importance and economic value, the family Lutjanidae (snappers) provides a compelling system for investigating connectivity processes. Snappers are key contributors to the global blue economy (FAO, 2022) and are ecologically important mid- to upper-trophic predators that influence prey populations, structure food webs, and support biodiversity (Szedlmayer & Lee, 2004; Wares & Szedlmayer, 2016). Red Snapper (*Lutjanus campechanus*) and Vermilion Snapper (*Rhomboplites aurorubens*) support important commercial and recreational fisheries across the northern Gulf of America (previously named “Gulf of Mexico” and hereafter referred to as “Gulf”), including the banks sampled in this study (Figure 1; Carpenter, 1965; Hood et al., 2007; Porch et al., 2007; Rindone et al., 2015; Spencer & Bruno, 2019). Although both species co-occur broadly from Texas to Florida, they differ in life-history characteristics. Red Snapper is a larger, longer-lived demersal species that undergoes ontogenetic habitat shifts, whereas Vermilion Snapper is generally smaller, shorter-lived, and earlier-maturing, typically occurring in schools over shelf-edge and hard-bottom habitats (Grimes, 1978; Grimes & Huntsman, 1980; Wilson & Nieland, 2001; Fischer et al., 2004; SEDAR, 2016; SEDAR, 2018; SEDAR, 2024; Moncrief et al., 2018; Chamberlin et al., 2023). Despite differences in life-history characteristics, each species is managed as a single Gulf-wide stock.

**Figure 1.**
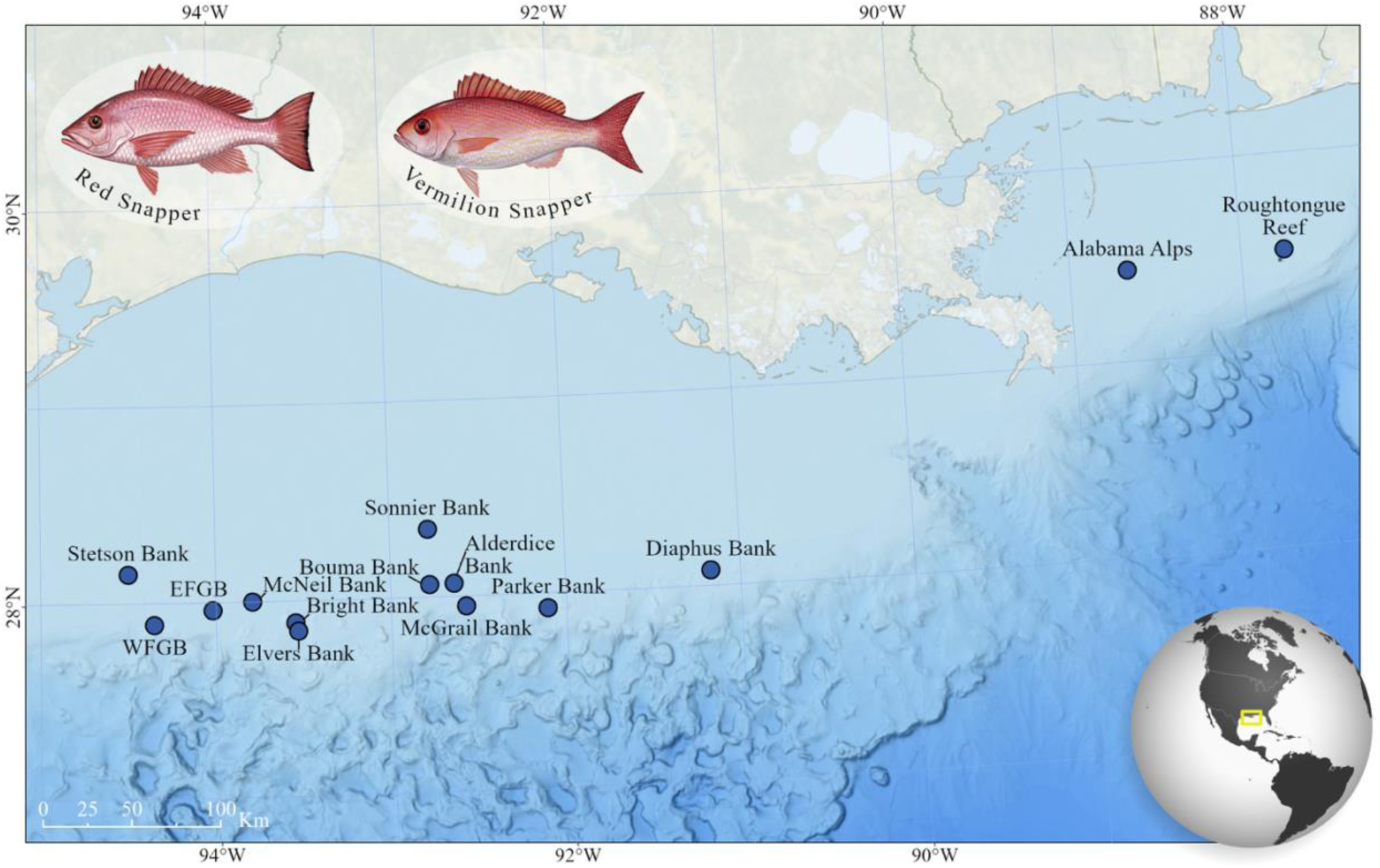
Map of the shelf-edge banks sampled for Red Snapper and Vermilion Snapper. Sites indicate locations where both otolith and genomic samples were collected. Snapper illustrations courtesy of NOAA Fisheries. Basemap data courtesy of the General Bathymetric Chart of the Oceans and NOAA’s National Centers for Environmental Information.

For Red Snapper, decades of molecular and genomic studies have identified high connectivity and weak population structure across much of the Gulf of American Basin (Gold et al., 1997, 2001; Pruett et al., 2005; Saillant & Gold, 2006; Hollenbeck et al., 2015; Portnoy et al., 2022). However, pronounced spatial variation in recruitment, growth, and fishing pressure, along with rebuilding trajectories, has led to more spatially explicit assessment models while retaining a single biological stock definition (Fischer et al. 2004; Woods et al. 2003; Saari et al. 2014; Cass-Calay et al., 2015; SEDAR, 2016; SEDAR, 2024; Moncrief et al. 2018). In Vermilion Snapper, genetic connectivity remains unknown, yet the species is also managed as a single Gulf stock based on age-structured assessment models that account for uncertainty in spatial structure and demographic variation but do not currently support regional subdivision (SEDAR, 2016). Taken together, this management context highlights the need for integrative approaches that explicitly link connectivity, habitat use, and demographic processes to better understand population resilience and to inform sustainable management.

Among the various approaches used to assess connectivity, genomics and otolith microchemistry have become especially powerful tools for investigating habitat connectivity in commercially important marine species. Restriction site-associated DNA sequencing (RADSeq) generates thousands of genome-wide single nucleotide polymorphism (SNPs), providing fine-scale resolution of population structure and gene flow (Davey et al., 2011; Andrews et al., 2016). By enabling the discovery and genotyping of large numbers of SNPs across the genome, RADSeq provides greater power to detect subtle genetic differentiation in marine species, particularly with weak population structure (Benestan et al., 2015, 2021; McKeown et al., 2020). Otolith microchemistry is complementary to genomic data and provides a chronological record of environmental exposure that indicates movements across habitats throughout an individual’s life (Campana & Thorrold 2001; Tanner et al., 2016). Otoliths are metabolically inert, calcium carbonate structures located in the fish inner ears that incorporate trace elements from surrounding water and diet as they grow (Campana et al., 1995; Campana et al., 2000; Campana & Thorrold, 2001; Sluis et al., 2012, 2015; Patterson et al., 2014) and can be used to determine their age (Vanderkooy et al., 2020). Laser Ablation–Inductively Coupled Plasma–Mass Spectrometry (LA-ICP-MS) allows quantification of elemental composition across otolith growth bands to reconstruct connectivity, ontogenetic habitat shifts, nursery habitats, and stock structure across the life span (Thorrold et al., 2001; Sluis et al., 2012; Sturrock et al., 2012; Aschenbrenner et al., 2016; Sih et al., 2022). Together, these methods enable the evaluation of connectivity across evolutionary, ecological, and ontogenetic timescales, yielding a more comprehensive understanding of population structure and habitat use.

To better understand habitat connectivity and demographic variation across snappers in the northern Gulf, we integrated genome-wide SNP data generated by RADSeq with otolith microchemistry to evaluate population connectivity and habitat use in two commercially important snapper species across the shelf-edge banks in the northwestern Gulf and interpret these patterns in the context of observed demographic and growth variation. This study forms part of the Connectivity of Coral Ecosystems (CyCLE) in the northwestern Gulf of Mexico, a multidisciplinary effort designed to quantify three-dimensional (3D) connectivity among shallow and mesophotic coral ecosystems in the northwestern Gulf (NOAA, 2019). Because CyCLE was developed to provide connectivity information and tools for resource managers, this framework supports evaluation of the expanded Flower Garden Banks National Marine Sanctuary (FGBNMS) and informs future marine protected area planning, adaptive management, and conservation of reef-associated habitats (Office of National Marine Sanctuaries, 2024).

Here, we examine regional connectivity across ecological and evolutionary scales in Red and Vermilion Snapper collected from 14 shallow and mesophotic shelf-edge banks. Specifically, we assess whether genome-wide genetic variation reveals population structure across geographic and depth gradients, and whether otolith elemental signatures capture fine-scale, site-specific patterns of environmental exposure and habitat use that are not resolved by genetic markers alone. We then relate these complementary connectivity signals to spatial variation in age structure and growth dynamics to evaluate how demographic processes operate within a regionally connected system. Together, this framework links regional genomic connectivity with otolith-derived environmental signatures and demographic variation, providing a multidimensional perspective on connectivity that informs understanding of population resilience and spatial management in reef-associated fisheries.

## 2. Materials and Methods

### 2.1. Study Location and Sample Collection

To investigate the processes shaping connectivity, Red Snapper (*Lutjanus campechanus*) and Vermilion Snapper (*Rhomboplites aurorubens*) were collected from 14 shelf-edge banks within the FGBNMS and neighboring areas of the northwestern Gulf: Stetson Bank, West Flower Garden Bank (WFGB), East Flower Garden Bank (EFGB), McGrail Bank, Bright Bank, Elvers Bank, McCall Bank, Bouma Bank, Sonnier Bank, Parker Bank, Alderdice Bank, Diaphus Bank, Alabama Alps, and Roughtongue Reef. Red Snapper was collected across all 14 banks, whereas Vermilion Snapper was collected at 11 of the 14 banks, with no specimens collected from EFGB, Bright Bank, or Bouma Bank. Sampling spanned shallow (15-40 m), upper mesophotic (40-85 m), and lower mesophotic (85-150 m) depth zones using standardized hook-and-line fishing methods (Figure 1). Collections were conducted in 2019, 2021, and 2022 aboard the R/V *Southern Journey*. For each individual, total length (TL), standard length (SL), and weight were measured, except during the 2022 expedition, when a scale was unavailable. Muscle and gill arch tissue samples were obtained from each individual, stored in 95% ethanol or flash-frozen, and then stored at −80 °C at the Laboratory of Analytical Biology, Smithsonian Institution National Museum of Natural History. Sagittal otoliths (left and right) were removed from each individual sampled and stored in dry envelopes for elemental analysis.

### 2.2. Population Genomics

#### 2.2.1. Molecular Laboratory Methods

Genomic DNA was isolated from muscle tissue using a DNeasy Qiagen (Venlo, Netherlands) tissue genomic DNA extraction kit and the AutoGenPrep 965 automated DNA extraction robot (AutoGen, Holliston, MA, USA). DNA concentration was quantified using the Quanti-IT dsDNA Broad Range Assay kit (Thermo Fisher Scientific, Waltham, MA, USA) on a Qubit (Lytchett Matravers, United Kingdom) 4.0 Fluorometer. DNA integrity was assessed using 2.0% agarose gel electrophoresis. Normalized (20 ng per ul), high-quality (230/260 and 260/280 ratios >1.8) DNA was sent to Floragenex, Inc. (Eugene, OR, USA) for RAD-Seq library preparation. Libraries were constructed for each of the 484 samples using the *SbfI* restriction enzyme. In total, 484 libraries (241 Red Snapper and 243 Vermilion Snapper) were sequenced on an Illumina (San Diego, CA, USA) NovaSeq6000 (100 bp single end; University of Oregon’s Genomics and Cell Characterization Core Facility).

#### 2.2.2. Bioinformatics Processing and SNP Filtering

Raw data were quality checked using the program FASTQC (Andrews 2010) and demultiplexed using process_radtags in STACKS v2.68 (Catchen et al., 2013) with the restriction enzyme sbfI and inline barcodes (-e sbfI --inline_null). Default cleaning parameters were used to remove reads with uncalled bases (-c) and discard reads in which the average quality score dropped below a Phred score of 10 whiting the default sliding window (-q), corresponding to approximately 90% base-call accuracy. RADseq data for each species were processed with iPYRAD v 0.9.97 (Eaton & Overcast 2020) using the Vermilion Snapper chromosome-level genome (Roa-Varón et al., 2025) as a reference (DDBJ/ENA/GenBank accession JBFPGM000000000). For each species, reads were mapped to the reference genome, and RAD loci were assembled from mapped reads using iPYRAD with default parameters, except for sequence similarity (clust_threshold = 0.85), max heterozygotes in consensus (max_Hs_consens = 0.5), and maximum missing data of 15%. After individual-level filtering, samples with excessive missing data were removed, resulting in the exclusion of 48 Red Snapper and 33 Vermilion Snapper individuals and final dataset of 204 Red Snapper and 231 Vermilion Snapper individuals retained for downstream analyses (Table 1, Supplementary Table S1).

**Table 1.** Summary of sample numbers by species, analytical approach, and depth stratum. Values indicate the number of individuals retained Population Genomics and Connectivity.

| Approach | Depth | Red Snapper<br>(n=252; 14 sites) | Vermilion Snapper<br>(n=264; 11 sites) | Total<br>Analyzed |
| --- | --- | --- | --- | --- |
| RadSeq | Shallow | 15 | – | 15 |
|  | Upper Mesophotic | 131 | 159 | 290 |
|  | Lower Mesophotic | 58 | 71 | 129 |
|  | Total per species | 204 | 230 | 434 |
| Ablated Otolith | Shallow | 20 | – | 20 |
|  | Upper Mesophotic | 166 | 161 | 327 |
|  | Lower Mesophotic | 64 | 73 | 137 |
|  | Total per species | 250 | 234 | 484 |
| Aged Otolith | Shallow | 20 | – | 20 |
|  | Upper Mesophotic | 165 | 156 | 321 |
|  | Lower Mesophotic | 64 | 73 | 137 |
|  | Total per species | 249 | 229 | 478 |

The resulting datasets were further filtered using VCFtools (Danecek et al., 2011) to retain only bi-allelic SNPs present in at least 70% of individuals across all populations, with a minimum minor allele frequency of 0.01. These include SNPs with low or extremely high coverage (min-meanDP 10, max-meanDP 100), SNPs with low mapping quality, SNPs not in Hardy–Weinberg Equilibrium (HWE), and physically linked SNPs.

SNP genotypes were imported from STRUCTURE-format files (*.str) using the read.structure function in adegenet. The Red Snapper dataset comprised 204 individuals and 20,954 loci, while the Vermilion Snapper dataset included 231 individuals and 15,395 loci. Individuals were first assigned to sampling sites by defining a bank-level factor stored as a genind object and used as a primary grouping variable in downstream analyses. Alternative grouping schemes were created by re-importing the same genotype dataset and replacing site assignments with depth strata (Shallow, Upper Mesophotic, Lower Mesophotic), sampling year (2019–2022), and age class (Juvenile, Adult, Unknown) (Supplementary Table S2).

#### 2.2.3. Genetic Diversity and Differentiation

Diploid RADseq SNP genotypes were analyzed in R v4.5.3 (R Core Team, 2026) using a workflow that (i) imported SNP data as a genind object using the adegenet package (Jombart & Ahmed, 2011), (ii) assigned individuals to biologically meaningful grouping factors (site, depth, year, age), and (iii) quantified genetic diversity, differentiation, and population structure using complementary multivariate and assignment-based approaches.

The genind object was converted to a hierfstat data frame using genind2hierfstat. Summary statistics, including heterozygosity components and F-statistics, were calculated with basic.stats. Genetic distances among sites were estimated with genet.dist, and pairwise differentiation among sites was quantified using Nei’s pairwise F_ST_ (pairwise.neifst, diploid = TRUE). Within-site inbreeding coefficients F_IS_ were also recalculated to evaluate whether positive values were consistent across sites or could result from pooling effects, locus-specific patterns, or data-quality artifacts. To further assess these signals, we examined individual heterozygosity, locus-and individual-level missingness, and kinship relationships, and sensitivity analyses were conducted under increasingly stringent filtering thresholds for minor allele frequency, missingness, and HWE.

#### 2.2.4. Multivariate and Bayesian Analyses of Genomic Structure

##### Principal Component Analyses (PCA)

To visualize genome-wide patterns of variation, genotype tables were generated from the genind object using tab (NA.method="mean") and analyzed with dudi.pca (ade4) (Dray & Dufour 2007), retaining three axes. Eigenvalue and variance summaries were visualized with fviz_eig, and individual scores were plotted with fviz_pca_ind in factoextra (Kassambara & Mundt 2020). Points were colored by the grouping factor (species, sites, depth, year, or age) using a color-blind-friendly viridis “plasma” palette (Garnier et al. 2023 Hollenbeck et al., 2015, Rocchini et al., 2024).

##### Discriminant Analyses of Principal Components (DAPC)

To complement PCA with a supervised, group-aware method, DAPC was performed with dapc in adegenet. For Red Snapper, Diaphus Bank and Elvers Bank were excluded from the site-based analysis prior to DAPC because each was represented by a single individual (Supplementary Table S1), potentially leading to unstable group parameterization. The number of retained PCs was optimized with optim.a.score to balance discrimination and overfitting, and the final DAPC was run using the optimized number of PCs and the appropriate number of discriminant functions (K-1). Results were visualized using DAPC scatter and density plots, depending on discriminant dimensionality, and with compoplot to display individual assignment probabilities across groups. The same framework was repeated for depth and year groupings to evaluate whether genetic structure aligned more strongly with geography, vertical habitat, or temporal sampling.

##### Bayesian Clustering Analyses

Population structure was inferred using the Bayesian clustering algorithm implemented in STRUCTURE, run through the command-line Python wrapper STRAuto (Chhatre & Emerson, 2017). Analyses evaluated a range of putative numbers of genetic clusters from K=1 to 14 for Red Snapper and from K=1 to 11 for Vermilion Snapper. For each *K*, 10 independent replicate runs were performed using a burn-in period of 100,000 steps followed by 250,000 Markov chain Monte Carlo (MCMC) iterations. Support for clustering was assessed across K values using mean log-likelihoods and consistency among replicate runs, and outputs were summarized and visualized using CLUMPAK (Kopelman et al., 2015).

### 2.3. Otolith Microchemistry and Demographic Analyses

#### 2.3.1. Otolith Preparation, Age Estimation, and Elemental Quantification

Sagittal otoliths were processed for two complementary purposes: age estimation and elemental chemistry. After cleaning and embedding, otoliths were transversely sectioned through the primordium. Thin sections were used for age estimation from annulus counts and marginal edge development, while thicker sections were prepared for LA-ICP-MS at Woods Hole Oceanographic Institution. We collected data on 12 analytes: lithium (7Li), boron (11B), magnesium (24Mg), calcium (43Ca and 48Ca), manganese (55Mn), cobalt (59Co), copper (63Cu), strontium (86Sr and 88Sr), and barium (137Ba and 138Ba). Elemental concentrations were first calibrated using the NIST 612 glass standard, with MACS-3 monitored as an internal standard, and corrected for time-varying background signal before being standardized as element:CA ratios. Prior to multivariate analyses, we also evaluated collinearity among otolith elements using correlation-based methods, hierarchical clustering, and variance inflation factors to assess redundancy and determine whether variable reduction was necessary. Full details of otolith preparation, age-reading procedures, LA-ICP-MS settings, elemental standardization, and collinearity analyses are provided in Supplementary Otolith Materials and Methods.

#### 2.3.2. Age Structure and Growth Modeling

Age structure and growth dynamics were examined for each species using otolith-derived age estimates and standard length (SL, mm) measurements. To identify the most appropriate functional form describing size-at-age relationships, age–length data for each species were evaluated within a multi-model inference framework incorporating several biologically plausible growth models commonly applied to marine fishes, including the von Bertalanffy growth function (VBGF), the Gompertz (Laird–Gompertz) model, and the logistic growth model (Katsanevakis 2006). All candidate models were fitted using nonlinear least squares estimation, and relative model support was assessed using the Akaike Information Criterion corrected for small sample size (AICc). Differences in AICc values (ΔAICc) and Akaike weights were used to quantify the strength of evidence for each model and to identify the most parsimonious representation of growth dynamics (Akaike 1974).

Growth models were first fitted at the species level to characterize regional growth patterns across shelf-edge banks. After establishing species-specific growth dynamics, demographic variation was evaluated by summarizing age distributions across sampling sites and visualizing them with boxplots. All analyses were conducted in R using base statistical functions and established growth-modeling workflows. This approach enabled direct comparison of growth trajectories and age structure between species while minimizing assumptions about site-specific differences in growth.

#### 2.3.3. Multivariate Analyses of Otolith Elemental Signatures

Multivariate analyses were conducted to evaluate spatial and biological drivers of otolith elemental composition. We first analyzed the combined dataset to test the effects of species, sampling site, and depth. We then performed species-specific analyses for Red Snapper and Vermilion Snapper to assess spatial structure among offshore banks and identify elemental gradients associated with these patterns. All multivariate analyses were conducted in R v4.5.3 (R Core Team 2026) using individual-level averages of elemental (Me:Ca) ratios derived from sagittal otoliths. Elemental variables were centered and scaled to unit variance prior to analysis to account for differences in magnitude among elements. Analyses were performed using the package vegan v 2.7.2 (Oksanen et al., 2025), while data manipulation and visualization were conducted using dplyr (Wickham et al., 2026), ggplot2 (Wickham 2016), ggrepel (Slowikowski 2024), viridis (Garnier et al., 2023), and patchwork (Pedersen 2024). Two Red Snapper sites with low sample sizes (Elvers, n = 1; Diaphus, n = 2) were excluded from site-based PERMANOVA, PERMDISP, pairwise site comparisons, and CAP analyses constrained by site due to insufficient replication. These samples were retained in depth-based analyses.

##### Principal Component Analysis (PCA)

To explore patterns in multivariate elemental composition, PCA was performed on scaled elemental data for the full dataset and for each species separately. PCA ordinations, biplots, and loading scores were used to identify the elements contributing most strongly to major axes of variation and to assess overlap among species, sites, and depth strata.

##### Permutational Multivariate Analysis of Variance (PERMANOVA)

Formal tests of multivariate differences in elemental composition were conducted using PERMANOVA (Anderson 2001) implemented in vegan v 2.7.2 (Oksanen et al., 2025). Euclidean distance matrices were calculated from the scaled elemental dataset, and PERMANOVA models were run using 9,999 permutations. Analyses were performed for (1) the combined dataset including species, site, and depth as predictors and (2) within each species separately to test for spatial and depth-related structuring. For each model, marginal tests (by = "margin") were used to evaluate the unique contribution of each predictor while controlling for the others.

##### PERMDISP and Pairwise Comparisons

To assess whether PERMANOVA results could be influenced by differences in group dispersion, tests for homogeneity of multivariate dispersion (PERMDISP; Anderson 2005) were conducted using the betadisper function in vegan.

Permutation tests (9,999 permutations) were used to determine whether observed multivariate differences indicated shifts in group centroids or unequal within-group variability. When PERMANOVA indicated significant site-level differences, pairwise PERMANOVA comparisons were performed among sites to identify which locations differed significantly in multivariate chemical composition. Resulting p-values were adjusted for multiple comparisons using the Benjamini–Hochberg false discovery rate (FDR; Benjamini & Hochberg 1995).

##### Canonical Analyses of Principal Coordinates (CAP) and Elemental Vector Fitting

To visualize constrained multivariate structure and maximize discrimination among spatial groups, CAP was performed using the capscale function. CAP ordinations were generated for the combined dataset and for each species separately using Euclidean distances. To identify elements most strongly associated with the ordination axes, elemental vectors were fitted onto CAP ordinations using the envfit function. Statistical significance of vector correlations was evaluated using 9,999 permutations, with α = 0.05. For elemental vector fitting, p-values were corrected for multiple testing using the FDR. Only vectors with adjusted p-values ≤ 0.05 were retained for visualization.

## 3. Results

### 3.1. Dataset Summary

A total of 516 specimens were collected across the study region. After quality control, retained sample sizes varied across RADseq, otolith chemistry, and age datasets as a result of genomic sequencing depth and assembly metrics, as well as variation in otolith readability.

Individuals excluded from RADSeq analyses were retained for otolith analyses when suitable and when material was available. Final sample sizes by species and depth are summarized in Table 1 and Supplementary Table S1.

### 3.2. Population Genomics and Connectivity

#### 3.2.1. RADSeq Data Processing and SNP Datasets

Across all samples, 913.2 million raw reads were processed in Stacks, and 716.1 million (78.4%) were retained after demultiplexing and quality filtering. In total, 21.6% of reads were removed because barcodes were not recovered; the RAD cut site was not detected, or reads were low in quality. Red Snapper samples retained a mean of 1,629,952 ± 1,869,050 SD reads per individual and 19,445 ± 1,930 SD loci per individual, yielding a final dataset of 20,954 loci with a maximum of 15.2% missing data per locus. Vermilion Snapper samples retained a mean of 1,955,990 ± 1,773,295 SD reads per individual and 14,024 ± 1,793 SD loci per individual, yielding a final dataset of 15,395 loci overall, with a maximum of 14.7% missing data per locus.

#### 3.2.2. Population Structure, Genetic Differentiation, and Connectivity

Genetic diversity and differentiation statistics were calculated using sampling sites as predefined population units, with additional analyses grouping individuals by depth stratum, sampling year, and age classes. PCA, DAPC, and STRUCTURE were then used to evaluate whether these predefined groups corresponded to discrete genetic clusters or instead indicated regional connectivity.

Overall patterns of genetic diversity and differentiation were broadly similar between Red Snapper and Vermilion Snapper, although subtle differences were evident. Vermilion Snapper exhibited slightly higher genetic diversity, with greater observed heterozygosity (H_O_ = 0.053) and expected heterozygosity within and across sites (H_s_ and H_T_ = 0.076) compared to Red Snapper (H_O_ = 0.050; Hs and H_T_ = 0.067; Table 2). Despite these differences in diversity, both species showed low among-site differentiation. Mean fixation indices F_ST_ were near zero in both species, indicating little to no detectable genetic differentiation among sampled banks. Red Snapper showed slightly higher overall F_ST_ than Vermilion Snapper (F_ST_ = 0.007 vs 0.001; Table 2), but both values were very low and consistent with broad genetic connectivity and no evidence of biologically meaningful population subdivision at the scale examined.

**Table 2.**
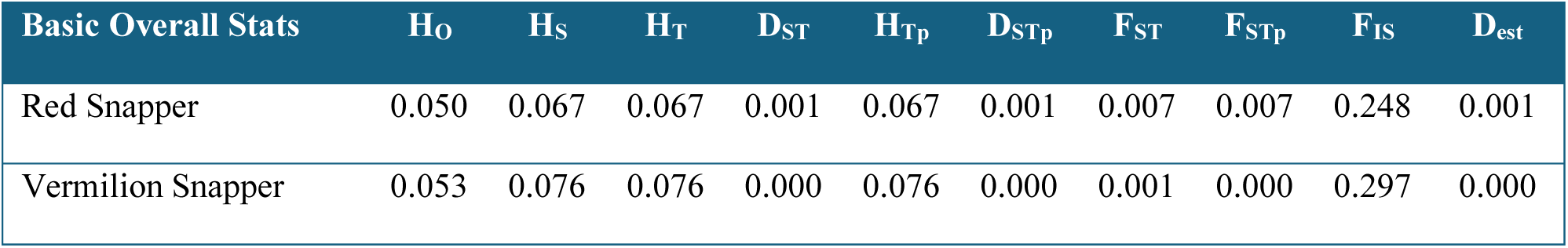
Overall genetic diversity and differentiation statistics per snapper species. Observed heterozygosity (H_O_), within-population gene diversity (H_S_), total gene diversity (H_T_), gene diversity among samples (D_ST_), corrected total gene diversity (H_tp_), gene diversity among samples (D_st_), corrected total gene diversity (H_Tp_), corrected gene diversity among samples (D_STp_), fixation index (F_ST_), corrected fixation index (F_STp_), inbreeding coefficient (F_IS_), Jost’s Differentiation estimator (D_est_).

Consistent with the F_ST_ results, gene diversity among samples (D_ST_), corrected gene diversity among samples (D_STp_), and Jost’s differentiation estimator (D_est_) were near zero in both species (Table 2), further indicating minimal genetic subdivision across sampling sites. Inbreeding coefficients (F_IS_) were moderately positive in both species (0.248 in Red Snapper and 0.297 in Vermilion Snapper), suggesting heterozygote deficits. Because this combination of near-zero F_ST_ and positive F_IS_ can indicate hidden substructure or data-related artifacts, we evaluated whether this pattern resulted from a Wahlund effect (an apparent heterozygote deficit caused by hidden population subdivision, Wahlund 1928) or instead arose from data-related artifacts, including pooled samples, genotyping bias, and missing data (Garnier-Géré & Chikhi 2013; De Meeûs 2018). As part of this evaluation, F_IS_ was recalculated within each sampling site for both species (Table S3).

Elevated within-site F_IS_ values were driven by a subset of loci and were restricted to a small number of sites, including Sonnier Bank in Red Snapper, and Diaphus, Alderdice, and Alabama Banks in Vermilion Snapper, while median locus-specific values remained near zero across most sites. In Red Snapper, elevated F_IS_ at Sonnier coincided with reduced observed heterozygosity and higher missingness. Kinship analyses did not detect close relatedness among individuals in either species. Sensitivity analyses using alternative filtering thresholds, including missingness thresholds of <15%, <20%, <25%, and <30%, and minor allele frequency (MAF) thresholds of >0.01 and >0.02, reduced the number of retained loci but did not reveal consistent genome-wide patterns of inbreeding across sites. Within-site F_IS_ was recalculated using a filtered dataset that retained loci with <30% missing data and MAF >0.01, yielding 5,085 loci for Red Snapper and 3,181 for Vermilion Snapper.

Pairwise Dch genetic distances among sampling sites were uniformly low for both species (Table S4a-b), consistent with weak spatial differentiation. In Red Snapper, pairwise Dch distances ranged from 0.010 to 0.050, with a mean distance of approximately 0.020. Vermilion Snapper exhibited consistently lower and more homogeneous genetic distances, ranging from 0.008 to 0.015, with a mean of approximately 0.012.

Population structure analyses supported a single broadly connected genetic unit for each species across the sampled banks. PCA and DAPC analyses revealed no discrete genetic clusters in either Red Snapper or Vermilion Snapper, and STRUCTURE was consistent with this pattern, supporting *K* = 1 for both species. Thus, although individuals were sampled across multiple banks, years, age classes, and depth strata, the genomic data did not support subdivision into multiple populations at the spatial scale examined.

Principal Component Analysis (PCA) revealed no discrete clusters within either species; rather, extensive overlap among individuals from different banks (Figure 2). The first two principal components explained a small proportion of the genetic variance (PC1 ∼ 1 %; PC2 ∼0.9 %), consistent with low among-group structure. Red Snapper exhibited slightly greater dispersion along PC1 relative to Vermilion Snapper, whereas Vermilion Snapper individuals were more tightly centered around the origin. However, in both species, ordinations failed to identify evidence of discrete subpopulations. Additional PCA projections stratified by age classes, sampling years, and depth strata, with no clear clustering along the major axes of genetic variation for Red Snapper (Supplementary Figure S1a-c) and Vermilion Snapper (Supplementary Figure S2a-c).

**Figure 2.**
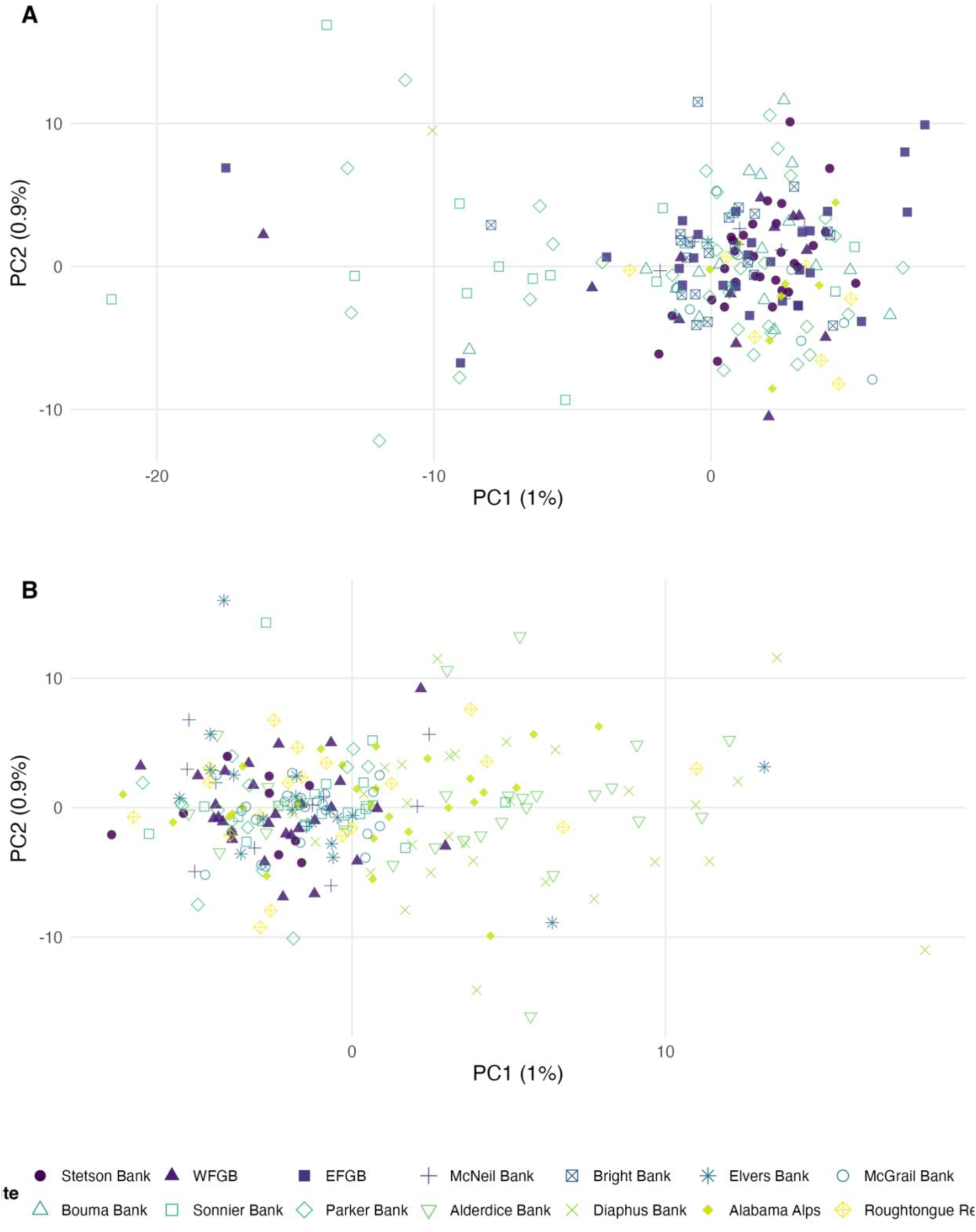
Principal components analysis of multilocus SNP genotypes across shelf-edge coral banks in the northern Gulf for (A) Red Snapper and (B) Vermilion Snapper. Individuals are colored and shaped by the sampling site. The first five PCs each explained < 1.5% of the variance in both species.

DAPC analyses similarly showed extensive overlap among predefined groups in both species, with no strong separation by site, depth stratum, sampling year, or age class. Depth-based DAPC showed broad overlap among shallow, upper, and lower mesophotic samples in both species (Figure 3A-B), indicating weak or undetectable depth-associated genetic structure. Site, year, and age-based DAPC analyses showed similar overlap, with no clear clustering by predefined groups in either species (Supplementary Figures S3–S4). STRUCTURE results were consistent with these multivariate analyses, with the strongest support for *K* = 1 in both species and no evidence of individual admixture patterns associated with site, depth, year, or age class.

**Figure 3.**
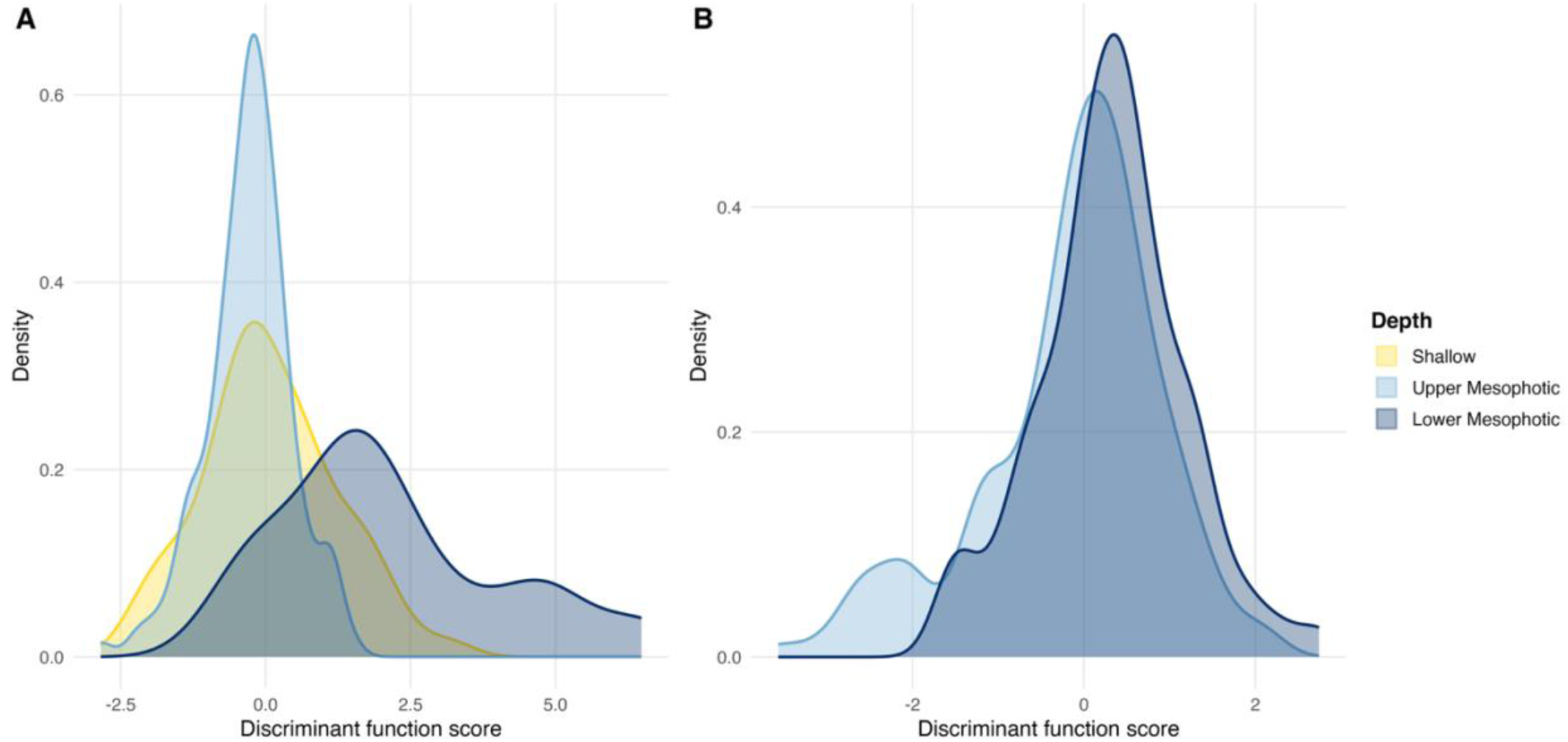
Density distributions of discriminant function scores from DAPC analyses by depth for (a) Red Snapper and (b) Vermilion Snapper. Shallow, upper mesophotic, and lower mesophotic groups show extensive overlap in discriminant space, indicating weak or undetected depth-associated genetic structure in both species.

### 3.3. Otolith-Derived Demographic and Environmental Structure

#### 3.3.1. Age and Growth

Comparative growth analyses revealed contrasting patterns of model support between Red Snapper and Vermilion Snapper when age–length relationships were evaluated within a multi-model inference framework. For Red Snapper, model selection based on AICc indicated that the logistic growth model received the strongest empirical support (Akaike weight = 0.52), with the Gompertz model also competitive (ΔAICc < 1), whereas the von Bertalanffy growth function (VBGF) received comparatively less support. In contrast, Vermilion Snapper exhibited substantial model uncertainty, with the VBGF, Gompertz, and logistic models receiving nearly identical support (ΔAICc ≤ 0.25; Akaike weights ≈ 0.31–0.35). Together, these results indicate species-specific differences in growth dynamics and underscore the importance of evaluating multiple growth models rather than relying on a single assumed functional form.

These species-specific growth patterns were accompanied by differences in demographic structure. Red Snapper spanned a wider age and size range, including individuals up to approximately 20 years, whereas Vermilion Snapper showed a more strongly young-skewed age distribution, with most individuals concentrated between 2 and 10 years, despite a few older individuals approaching 19 years. These differences were demonstrated in the growth models, with Red Snapper attaining a higher asymptotic length and continuing somatic growth later in life, while Vermilion Snapper approached a lower asymptotic size more rapidly (Figure 4).

**Figure 4.**
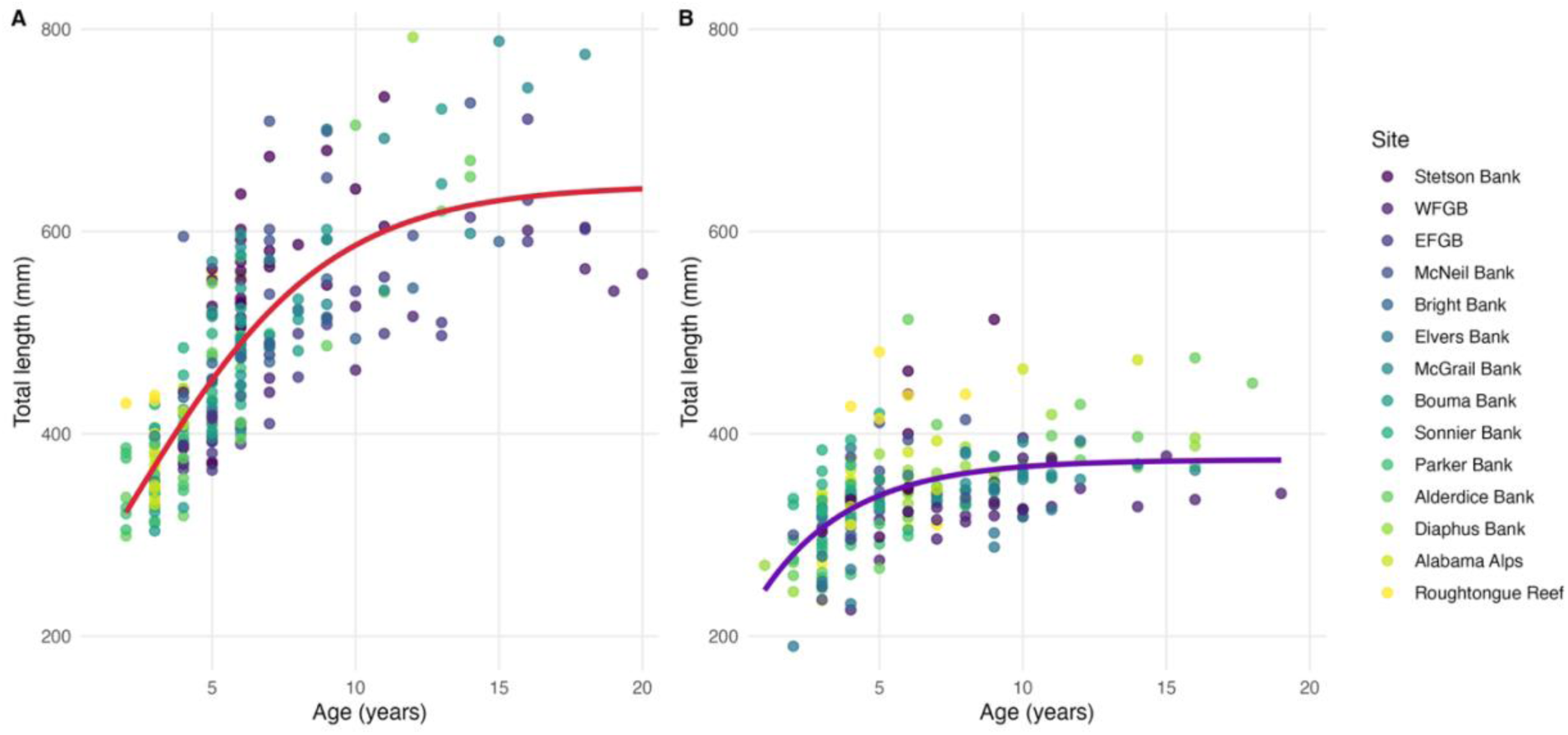
Species-level age-length relationships and fitted growth curves for (A) Red Snapper and (B) Vermilion Snapper. Points represent individual fish colored by sampling site. Solid lines show the fitted growth model with the strongest AICc support for each species.

Age structure varied among sites for both species (Figure 5A-B, Supplementary Table S5). Red Snapper exhibited a broad age distribution across shelf-edge and reef sites, ranging from 3 to 18 years. Older individuals (>15 years) were present at WFGB, EFGB, McGrail, whereas Sonnier, Parker, Alabama Alps, and Roughtongue Reef were dominated by younger coho rts; therefore, age structure varied among sites, with some banks containing mixed age classes and others composed primarily of younger individuals. Vermilion Snapper age distributions were narrower across shelf-edge banks and reef sites, with most individuals falling between 2 and 12 years.

**Figure 5.**
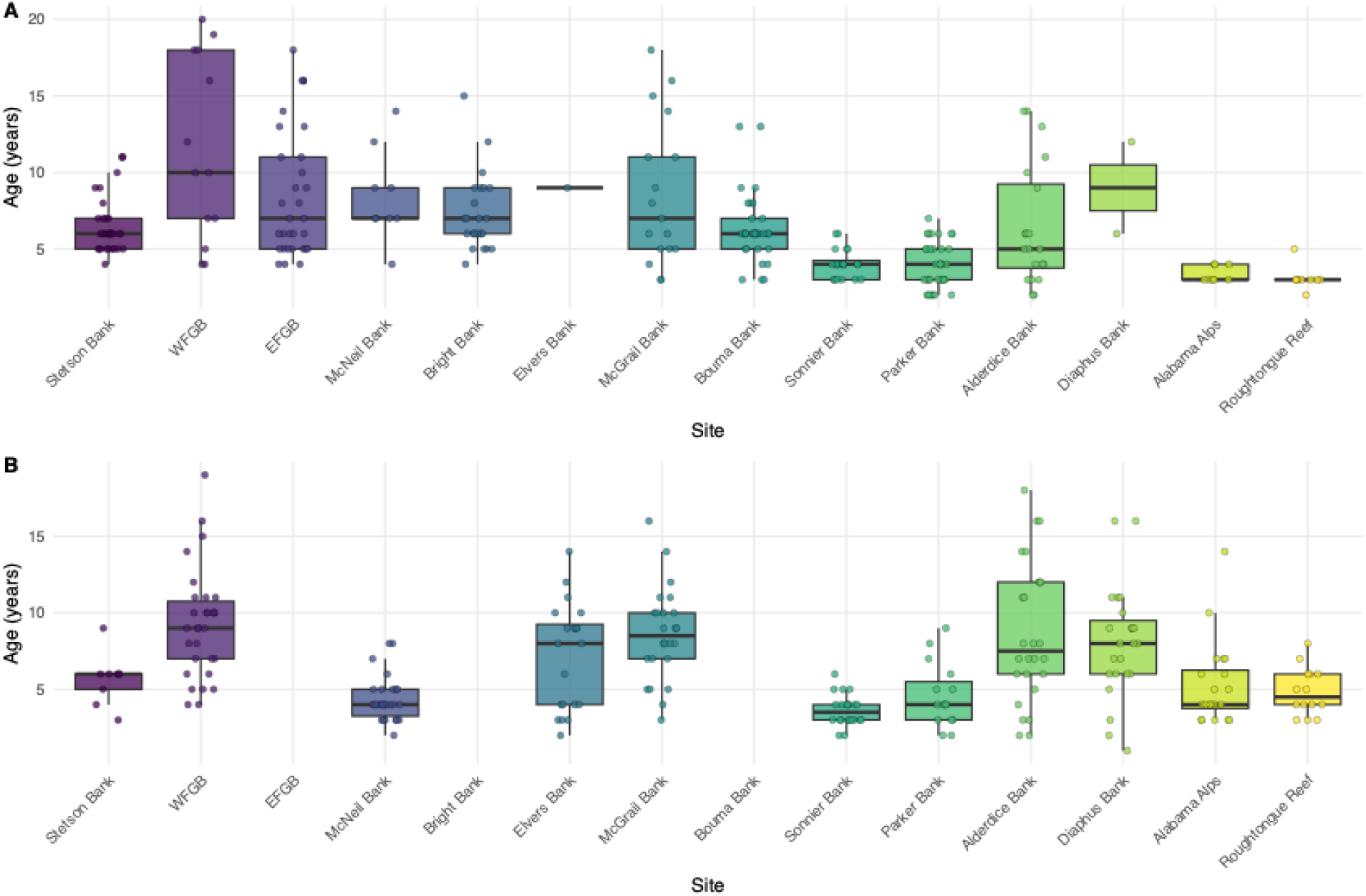
Age structure of (a) Red Snapper and (b) Vermilion Snapper across shelf-edge banks. Individuals are colored by sampling site, with site colors ordered according to geographic positions in Figure 1 and plotted from west to east. Boxes represent the interquartile range (25th–75th percentiles), horizontal lines indicate medians, and whiskers extend to the most extreme observations within 1.5 times the interquartile range. Points represent individual fish ages.

Older individuals (>12 years) occurred at WFGB, Alderdice, and Diaphus, whereas Stetson, Sonnier, and Alabama Alps, and Roughtongue Reef were dominated by younger cohorts.

Within-site age ranges were consistently narrower than those of Red Snapper, and maximum ages were lower across the sampled system.

#### 3.3.2. Collinearity Among Otolith Elemental Variables

In both species, most pairwise correlations among elemental ratios were low to moderate, indicating limited redundancy across the multielement dataset and no strong evidence that variation was dominated by shared geochemical controls (Supplementary Table S6, Supplementary Figure S5). For example, in Red Snapper, the highest correlations were limited to a small subset of elemental pairs, including B-Co, Co-Cu, and Li-Co. In Vermilion Snapper, correlations were lower overall, with moderate associations observed for Co-Cu, B-Co, and Li-Ba. Variance inflation factors were also generally low, with most VIF values below commonly used thresholds for moderate collinearity (VIF < 5), indicating that multicollinearity was unlikely to bias downstream multivariate analyses (Supplementary Table S7). Where elevated correlations or VIF values were observed, these were limited to a small number of elements and were not consistent across both species, indicating species-specific rather than systematic covariance. Together, these results support retaining the full elemental suite for subsequent multivariate analyses because the otolith chemistry dataset does not show strong multicollinearity in either Red Snapper or Vermilion Snapper.

#### 3.3.3. Otolith Elemental Signatures Across Species, Sites, and Depths

Unconstrained principal component analysis (PCA) of scaled otolith elemental ratios revealed multivariate structure in otolith chemistry across individuals (Figure 6). The two principal components explained a substantial proportion of the total variance in otolith microchemistry, with PC1 and PC2 accounting for 30% and 23% of the variance, respectively. In the combined ordination, samples showed partial separation by species, with Red Snapper tending to occupy more positive PC1 values and Vermilion Snapper distributed more broadly toward negative PC1 values, although overlap between species remained substantial (Figure 6A).

**Figure 6.**
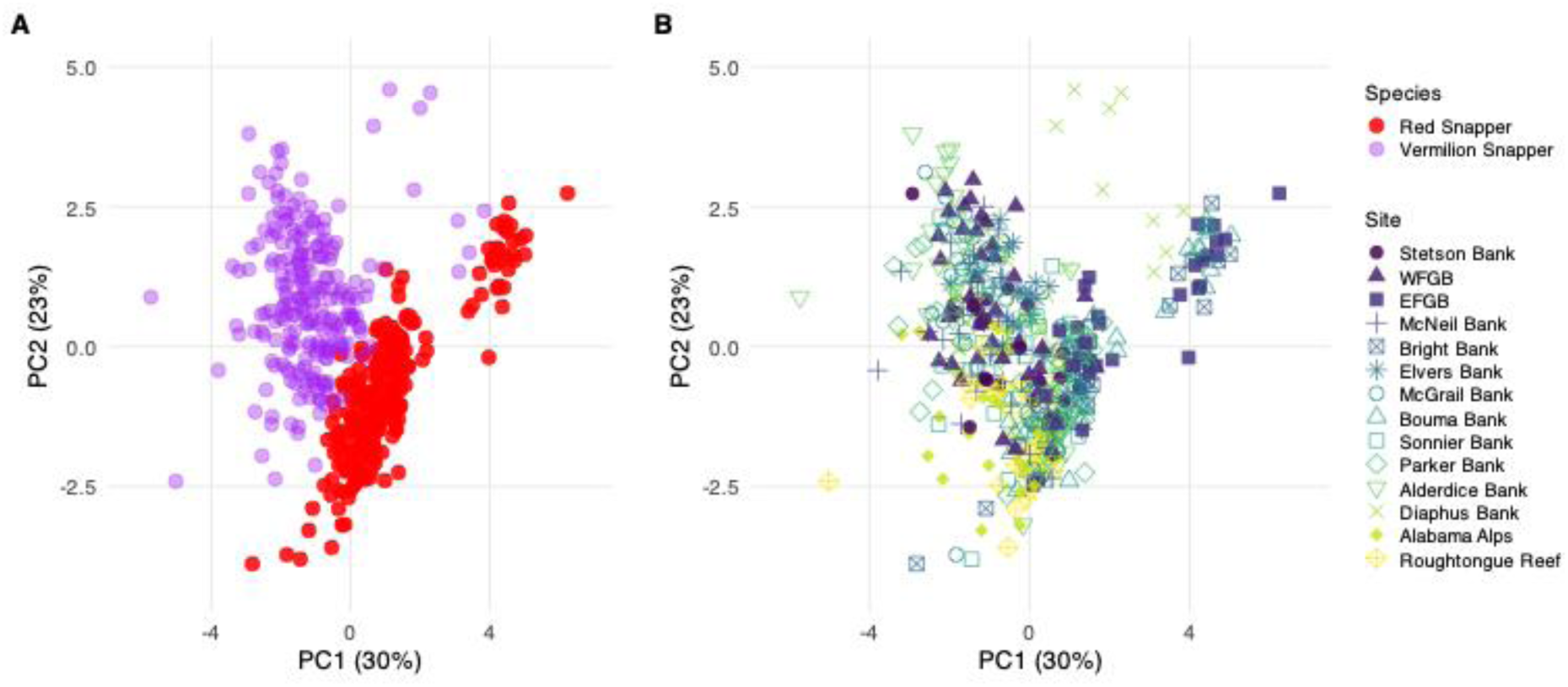
Principal component analysis (PCA) of scaled otolith elemental composition for Red Snapper and Vermilion Snapper across the northwestern Gulf. (A) Points represent individual fish colored by species. (B) Points represent individual fish colored by sampling site.

Otolith elemental composition showed bank-level spatial heterogeneity across the study region, despite broad overlap among sites (Figure 6B). In the site-colored ordination, samples from several offshore banks occupied partially distinct regions of multivariate space. This pattern was more apparent in Diaphus Bank, represented primarily by Vermilion Snapper, whereas samples from EFGB, Bright Bank, and Bouma Bank were represented only by Red Snapper. Although incomplete species representation at some banks limits direct site comparisons, the ordination suggests spatial variation in multielement otolith signatures among banks.

PCA ordinations examined by depth stratum and ontogenetic stage showed weaker structure than the species- and site-based ordinations. Although the first two axes captured a substantial proportion of the total otolith chemical variation (PC1 = 30%; PC2 = 23%; Figure 7), they did not separate individuals by depth or life stage. Within both species, individuals from shallow, upper mesophotic, and lower mesophotic habitats overlapped broadly in ordination space (Figure 7A). Likewise, juveniles and adults showed substantial overlap within each species (Figure 7B). These results indicate that depth stratum and ontogenetic stage were not the dominant sources of separation along the main axes and contributed less to the observed multivariate structure than species and sampling site.

**Figure 7.**
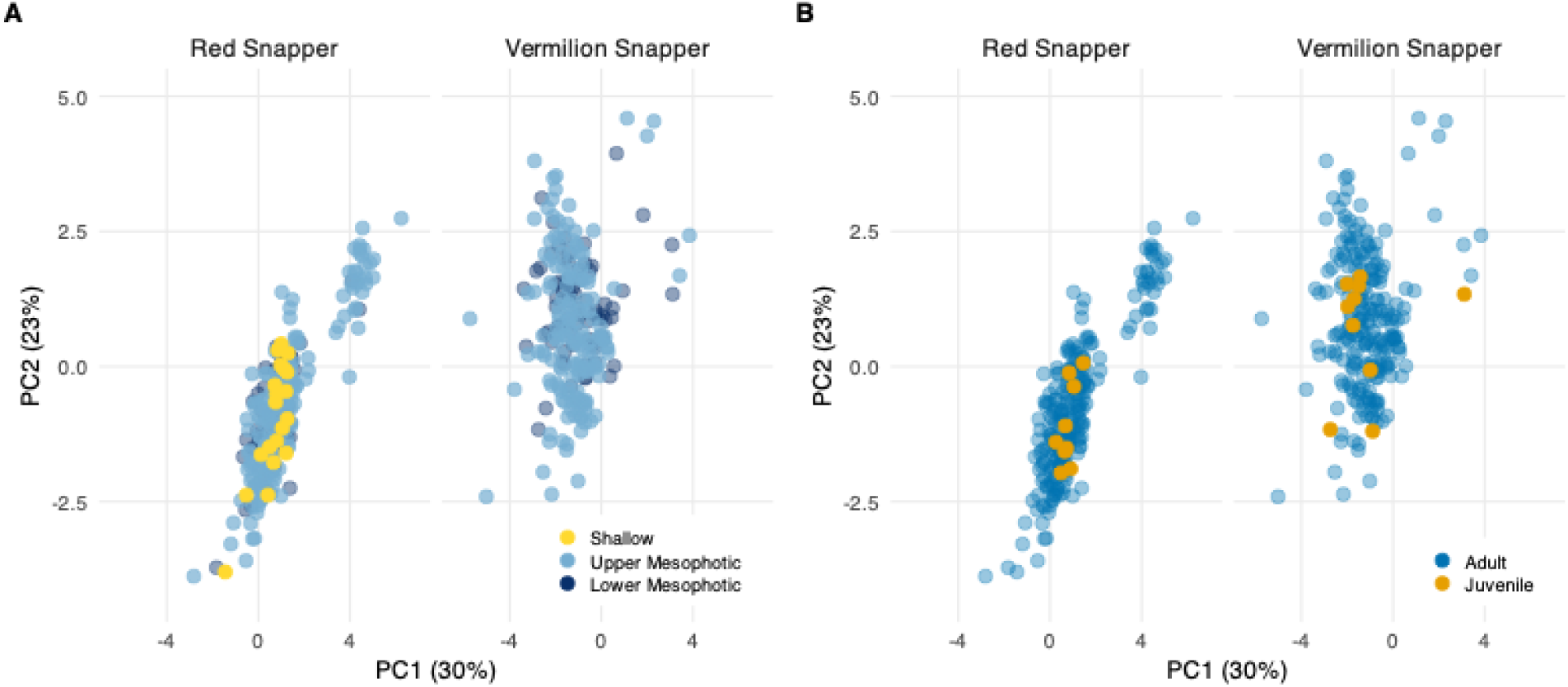
Principal component analysis (PCA) of scaled otolith elemental composition for Red Snapper and Vermilion Snapper. (A) Individuals colored by depth. (B) Individuals colored by age class.

The WFGB-focused PCA showed partial separation by species and depth, but substantial overlap among groups, indicating that these patterns should be interpreted cautiously (Supplementary Figure S6). WFGB was analyzed separately because it was the only site where both species were represented across upper and lower mesophotic strata with sufficient samples for qualitative comparison. Uneven sampling across sites and depth strata limits strong inference about depth-associated otolith differentiation.

#### 3.3.4. Spatial Structure and Elemental Drivers of Otolith Chemistry Combined-Species Structure

PERMANOVA results revealed significant effects of all three predictors (species, sites, and depth; p < 0.001), with sampling site explaining the largest proportion of variance (R² = 0.167), followed by species identity (R² = 0.121), whereas depth accounted for a comparatively small fraction of the total multivariate variation (R² = 0.014) (Table S8).

To assess whether PERMANOVA results could be influenced by differences in within-group dispersion, homogeneity of multivariate dispersion (PERMDISP) was evaluated for species, site, and depth. No significant differences in dispersion were detected between species (p = 0.894) or among depth strata (p = 0.103), indicating that differences associated with these predictors primarily resulted from shifts in group centroids. In contrast, dispersion differed significantly among sampling sites (F = 3.66; p < 0.001), indicating that site-level differences in otolith microchemistry may result from both shifts in centroid position and variation in within-site variation dispersion (Supplementary Table S8; Figure 8).

**Figure 8.**
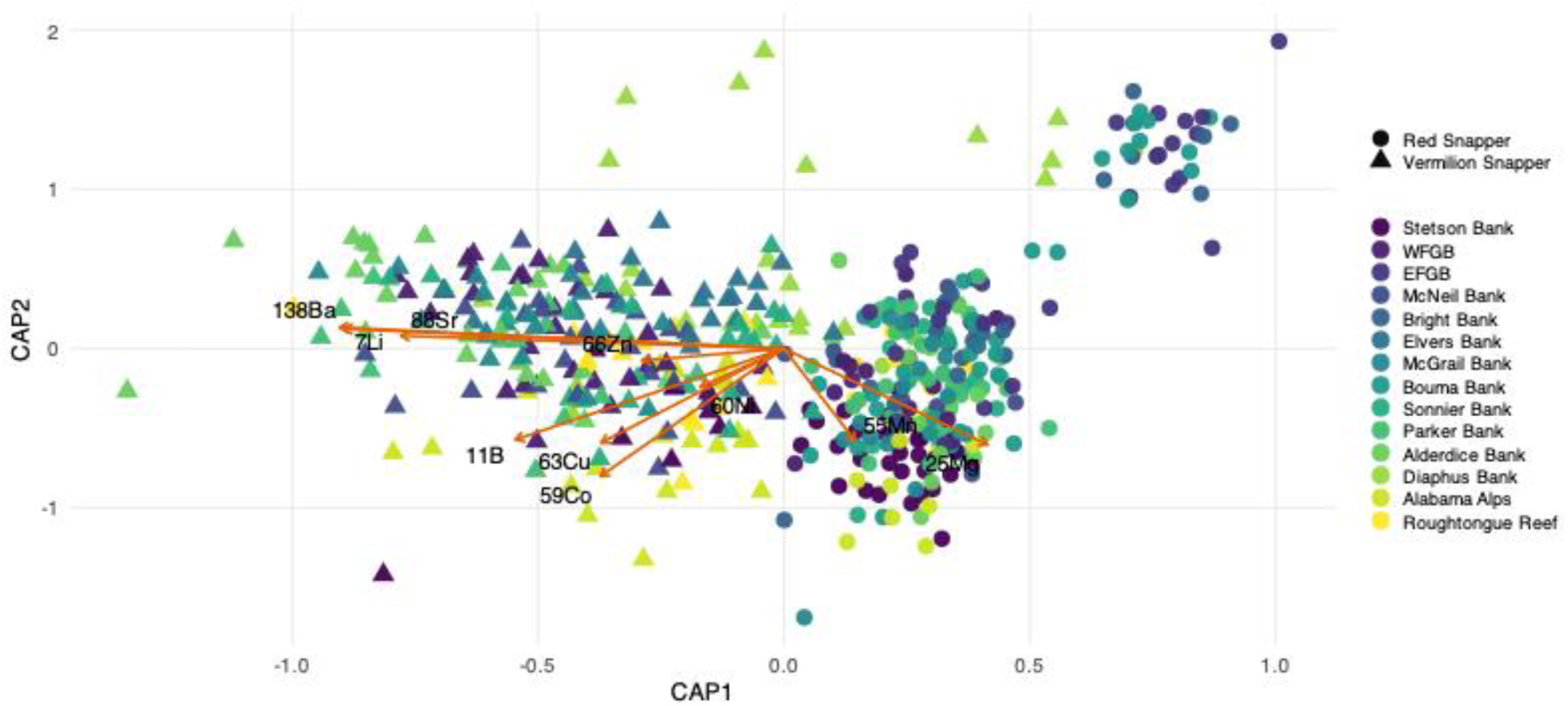
Canonical analysis of principal coordinates (CAP) of scaled otolith elemental composition constrained by sampling species, sampling site, and depth. Points represent individual fish and are colored by sampling site, with symbol shape indicating species. Vectors show the direction and relative contribution of elemental variables to the constrained axes.

To visualize the multivariate structure identified in PERMANOVA, a canonical analysis of principal coordinates (CAP) was performed using species, sampling sites, and depth as constrained variables. The overall CAP model was highly significant (F = 17.73, p < 0.001), indicating that these predictors collectively captured significant structure in otolith elemental composition. The CAP ordination showed partial separation between Red Snapper and Vermilion Snapper, with additional site-level structuring across the study region (Figure 8). Depth contributed relatively little to the overall constrained variation and is not shown as a separate visual grouping in the ordination.

All elemental vectors were significantly associated with the CAP ordination (permutation tests, p < 0.001). Ba showed the strongest fit to the ordination configuration (r² = 0.70), followed by Li (r² = 0.67), and Co (r² = 0.61), with B and Sr also showing strong fits. These results indicate that localized environmental conditions among banks are represented in multielement otolith fingerprints shared across both species, while species identity also contributes to the observed structure (Figure 8).

##### Within-Species Spatial Structure

To determine whether the spatial patterns detected in the combined analysis were consistent within each species, PERMANOVA analyses were conducted separately for Red Snapper and Vermilion Snapper using sampling site and depth as predictors (Supplementary Table S9).

Because these species-specific models were used as a follow-up analysis to the combined PERMANOVA, p-values were interpreted in relation to sample sizes and the consistency of patterns across models. Sampling site explained a substantial proportion of the multivariate variation (Red Snapper: R² = 0.225, F = 7.26; Vermilion Snapper: R² = 0.251, F = 7.48; p < 0.001). In contrast, depth-related variation differed between species. Depth had a statistically significant but weak effect in Red Snapper (R² = 0.016, F = 5.24, p < 0.001), whereas no significant depth effect in Vermilion Snapper (R² = 0.004, F = 1.21, p = 0.283). In both cases, horizontal variation among banks explained substantially more variation in otolith chemistry than depth strata.

Tests for homogeneity of multivariate dispersion (PERMDISP) indicated that dispersion differed significantly among sites in both species (Red Snapper: F = 3.31, p < 0.001; Vermilion Snapper: F = 2.76, p = 0.004), whereas depth-related dispersion was significant only in Red Snapper (F = 4.17, p = 0.016) and not in Vermilion Snapper (p = 0.676). These results indicate that site-level differences result from both centroid separation and within-site variability, while depth effects are weak and inconsistent across species. Pairwise PERMANOVA comparisons further resolved spatial structure in both species (Supplementary Table S9). In Red Snapper, most bank-to-bank comparisons were significant after multiple-testing correction, indicating widespread differentiation across the study region. Particularly strong contrasts involved EFGB relative to Roughtongue Reef and Parker Bank, as well as Stetson Bank versus EFGB, which showed among the highest levels of multivariate differentiation. In Vermilion Snapper, pairwise comparisons similarly revealed extensive spatial differentiation, with most banks differing significantly from one another. Strong differentiation was observed for comparisons involving Diaphus Bank, Alabama Alps, and Elvers Bank, which were consistently distinct from multiple other banks. However, a small number of non-significant comparisons (e.g., WFGB versus McGrail Bank) demonstrate partial overlap in otolith chemical signatures among some sites, indicating slightly weaker or more continuous spatial structuring relative to Red Snapper.

CAP ordinations constrained by site revealed concordant spatial gradients in both species, characterized by directional separation of banks along the primary axes rather than discrete clustering (Figure 9). These gradients broadly represented the geographic distribution of banks across the study region. In Red Snapper, banks such as EFGB and Bright Bank were positioned toward positive CAP1, whereas Alabama Alps and Roughtongue Reef were distributed among the easternmost sampled sites toward negative CAP1; the remaining banks occupied intermediate positions. Vermilion Snapper showed a similar pattern, with Diaphus Bank toward the positive end of CAP1 and Alabama Alps and Roughtongue Reef toward the opposite end of the ordination. These gradients indicate spatial variation in otolith elemental composition among banks but did not align with depth strata, which is consistent with the PERMANOVA results showing that sampling site explained more variation than depth.

**Figure 9.**
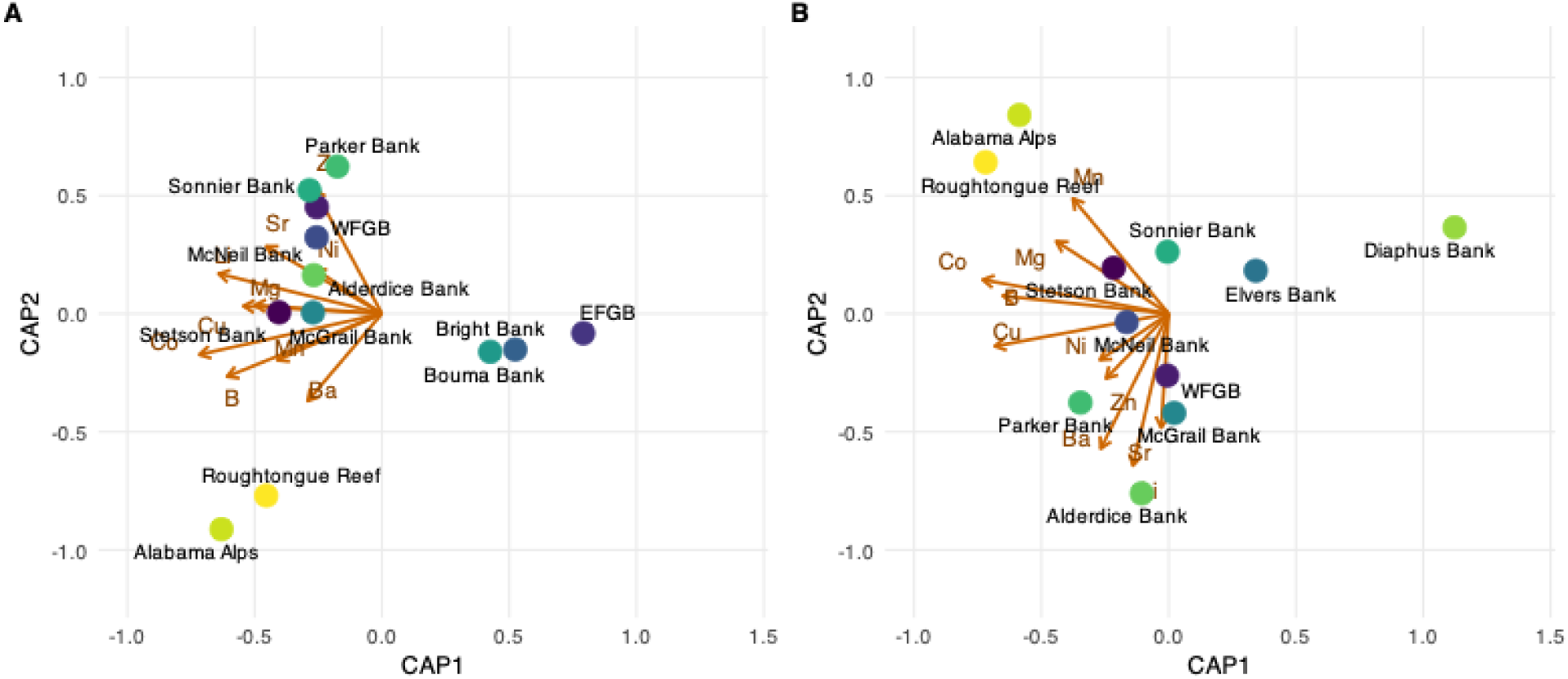
Canonical analysis of principal coordinates (CAP) centroid plots showing site-level multivariate structure in otolith elemental composition for (A) Red Snapper and (B) Vermilion Snapper across shelf-edge banks in the northern Gulf. Points represent site centroids and do not represent within-site dispersion; arrows indicate fitted elemental vectors.

Elemental vectors fitted onto the CAP ordinations indicated that spatial differentiation in both species is driven by multiple overlapping elemental signals. Across species, Co, Li, and B consistently exhibited strong associations with the constrained axes, with additional contributions from Cu, Ba, Mn, Sr, and Zn. These results indicate that localized environmental gradients among offshore banks are represented in multielement otolith fingerprints in both species, whereas depth-related gradients contributed relatively little to the observed structure.

## 4. Discussion

Otolith microchemistry revealed fine-scale habitat structure among shelf-edge banks that was not mirrored by genome-wide population structure. Although Red Snapper and Vermilion Snapper showed weak or undetectable genetic differentiation across the sampled region, otolith elemental composition varied among banks and between species, indicating localized environmental exposure of habitat use within a broadly connected genomic system. This contrast highlights the value of integrating markers that capture different biological scales: genome-wide SNPs demonstrate regional gene flow, whereas otolith chemistry provides an integrated record of environmental exposure across otolith growth.

### Otolith Microchemistry Reveals Fine-Scale Habitat Structuring Among Shelf-Edge Banks

Otolith elemental composition revealed fine-scale spatial structure among shelf-edge banks in both Red Snapper and Vermilion Snapper. This bank-level variation indicates that individuals experienced localized environmental conditions or habitat-use histories that were retained in the otolith chemical composition, even though sites showed broad overlap in multivariate space.

These results suggest that shelf-edge banks are not chemically or ecologically equivalent habitats, consistent with the ability of otolith microchemistry to resolve spatial variation in environmental exposure and habitat use (Campana 1999; Campana & Thorrold 2001; Elsdon et al., 2008; Tanner et al., 2016). When interpreted alongside genomic results, this otolith variation indicates spatial heterogeneity not indicated in genome-wide population structure, highlighting how geochemical and genetic approaches can complement each other when assessing connectivity (Standish et al., 2008; McKeown et al., 2015; Bertram et al., 2022).

Because otolith elemental values were averaged from core to edge, these patterns should be interpreted as integrated signals of environmental exposure across otolith growth rather than stage-specific reconstructions of larval, juvenile, or adult habitat use. Early-life stage environmental differences may have contributed to the observed chemical fingerprints, particularly if individuals experienced different larval or settlement environments. However, because much of the otolith material in adult fish represents later juvenile and adult growth, early-life variation alone is unlikely to fully explain the extent of among-bank differentiation observed in adult individuals (Thorrold et al., 2001; Sturrock et al., 2012; Sih et al., 2022).

Similar decoupling between genetic connectivity and otolith-based environmental signatures has been reported in other reef-associated fishes, underscoring how genomic and otolith approaches are complementary for resolving processes operating across different spatial and temporal scales (Standish et al., 2008; McKeown et al., 2015; Bertram et al., 2022). In this context, the otolith results provide evidence that localized environmental exposure and habitat use can vary among banks, even within a regionally connected system.

### Species-Specific Otolith Signatures Suggest Different Habitat-Use Histories

Species-specific elemental patterns suggest that Red Snapper and Vermilion Snapper recorded localized habitat variation in different ways. In Red Snapper, multivariate differentiation was driven primarily by B, Co, and Mg, elements whose incorporation into otoliths can be influenced by local water chemistry, carbonate conditions, physiological regulation, and growth or calcification processes (Campana, 1999; Campana & Thorrold, 2001; Patterson et al., 2001a; Miller, 2009; Patterson et al., 2014; Limburg & Casini, 2019). These patterns are compatible with previous evidence that Red Snapper can exhibit reef-associated residency, moderate to high site fidelity, and relatively small home ranges after settlement (Patterson et al., 2001b; Topping & Szedlmayer, 2011; Piraino & Szedlmayer, 2014; Williams-Grove & Szedlmayer, 2016; Everett et al., 2020; SEDAR, 2024).

In contrast, Vermilion Snapper exhibited a broader and more complex elemental fingerprint, with strong contributions from Li, Mn, Cu, Ba, Ca, and Sr. These elements are often associated with variation in water-mass characteristics, salinity, sediment-water interactions, redox conditions, diet, growth, and physiological regulation. Lithium and Ba have been linked to distinct geochemical regimes, freshwater influence, upwelling, or water-mass variation, while Sr and Ca can reflect differences in salinity, water chemistry, and calcification dynamics (Campana, 1999; Bath et al., 2000; Secor & Rooker, 2000; Campana & Thorrold, 2001; Elsdon & Gillanders, 2003). More variable Mn may indicate exposure to changing water chemistry, hypoxic or reducing conditions, or sediment-water redox dynamics, whereas Cu may indicate dietary, physiological, or growth-related pathways (Limburg et al., 2011, 2015; Lindberg and Casini, 2018; Hüssy et al., 2021, 2024). Together, these patterns indicate that Vermilion Snapper experienced a more heterogeneous range of environmental conditions or habitat-use histories than Red Snapper.

These species-level differences are biologically meaningful because they indicate that co-distributed snappers can experience the same shelf-edge seascape in different ways. Red Snapper chemistry is consistent with stronger reef-associated residency or localized exposure, whereas vermilion Snapper seems to record broader environmental heterogeneity. Thus, the otolith results indicate generic among-bank variation and suggest species-specific ecological pathways through which local habitat structure is expressed within a genetically connected system.

Depth-related differences in otolith chemistry were weaker than horizontal variation among banks. Although Red Snapper showed weak depth-associated variation, this pattern should not be interpreted as evidence for an ontogenetic depth shift because age, site, and depth stratum were not fully independent in sampling design. Most banks were sampled within a single depth stratum, complicating efforts to separate depth effects from bank-specific environmental conditions or local age structure. In Vermilion Snapper, depth was not a significant predictor of otolith chemistry. Overall, these results indicate that bank-specific environmental conditions, rather than depth strata alone, were the primary drivers of otolith elemental composition across the sampled shelf-edge habitats.

### Age and Growth provide Demographic Context for Local Habitat variation

Age structure and growth patterns provided additional evidence that demographic variation can occur among banks within a regionally connected system. Both species were dominated by younger individuals, with most Red Snapper and Vermilion Snapper younger than 10 years of age. This youth-skewed age structure is consistent with patterns often observed in exploited reef-fish populations, including prior evidence for reduced representation of older Red Snapper in Gulf age-composition data and recent stock assessment results (Saari et al., 2014; Powers et al., 2018; SEDAR, 2024). More broadly, fishing can reduce the proportion of older individuals in exploited marine populations, often resulting in age-structure truncation and reduced demographic stability (Barnett et al., 2017; Charbonneau et al., 2022). However, because observed age composition can be affected by gear selectivity, sampling design, and habitat-specific occupancy of older individuals, the young-skewed age structure observed should be interpreted as consistent with, but not diagnostic of, exploitation-driven age truncation.

The demographic implications of this age structure likely differ between species because Red Snapper and Vermilion Snapper differ in longevity, growth, and reproductive biology. Red Snapper are long-lived, reach larger sizes, and show strong age-and size-dependent increases in reproductive output; therefore, reduced representation of older individuals may have greater implications for reproductive potential compared to Vermilion Snapper, particularly because peak reproductive contribution occurs later in life (Gallaway et al., 2020; SEDAR, 2018; SEDAR, 2024). In contrast, Vermilion Snapper mature earlier and spawn across a broader range of younger ages compared to Red Snapper, indicating that a similar young-skewed age distribution may not have equivalent demographic consequences (Moncrief et al., 2018).

Growth patterns further support species-specific demographic differences within the shelf-edge habitat network. Red Snapper exhibited a growth pattern consistent with continued somatic growth later in life, whereas Vermilion Snapper showed a more constrained size-at-age pattern. These differences indicate that local variation in habitat quality, prey availability, density dependence, depth-associated conditions, or fishing exposure may influence demographic trajectories after settlement, even when populations are regionally connected (Dance & Rooker 2019; Erisman et al., 2020; Goldstein & Sponaugle 2020; Streich et al., 2017). Together, the age and growth results provide demographic context for the otolith patterns, reinforcing that genetically connected banks may still differ in local ecological conditions relevant to growth and resilience.

### Genome-Wide SNPs Reveal Regional Genetic Connectivity Across Shelf-edge Banks

Genome-wide SNPs showed weak or undetectable population structure across shelf-edge bank in both species, providing an important contrast to the otolith results. Low differentiation metrics, extensive overlap in PCA and DAPC, and STRUCTURE support for K = 1 indicate that Red Snapper and Vermilion Snapper each form a broadly connected genomic unit at the spatial scale examined. Thus, the bank-level otolith differences are unlikely to indicate discrete population subdivision and instead point to localized environmental exposure, habitat use, or post-settlement processes within a genetically connected system.

The genomic pattern is consistent with previous work on Red Snapper, which has generally reported high connectivity and weak genetic structure across much of the Gulf, although subtle geographic differentiation can emerge at broader spatial scales, particularly near the West Florida Shelf (WFS) (Gold et al., 1997, 2001; Pruett et al., 2005; Saillant & Gold, 2006; Hollenbeck et al., 2015; Portnoy et al., 2022). Our sampling did not include WFS sites; therefore, we could not evaluate the previously reported regional signal. For Vermilion Snapper, available connectivity information remains more limited, but previous microsatellite work similarly indicated no evidence of discrete Gulf stocks, with weak isolation-by-distance across the broader Atlantic-Gulf range (Tringali & Higham, 2007). These patterns support high regional genomic connectivity across the sampled shelf-banks.

High genomic connectivity does not necessarily imply ecological or demographic uniformity among banks. In the Gulf, larval transport and post-settlement connectivity are shaped by the regional circulation, including the Loop Current, mesoscale eddies, local retention, larval behavior, and settlement processes. These mechanisms can generate spatially structured recruitment and demographic heterogeneity even when gene flow is sufficient to homogenize genome-wide variation (Karnauskas et al., 2017, 2022; Vaz et al., 2023; Zhou et al., 2024). In this context, our results support a dual pattern in which regional larval exchange may maintain connectivity, while bank-specific habitat conditions and post-settlement processes generate ecological variation recorded in otolith microchemistry. This contrast helps to reconcile long-standing debates regarding snapper population structure by showing that regional genetic connectivity can coexist with localized structuring.

## 5. Conclusions and Management Implications

Genome-wide SNPs and otolith microchemistry revealed different but complementary dimensions of connectivity in Red Snapper and Vermilion Snapper across shelf-edge banks in the northern Gulf. Genome-wide analyses indicated weak or undetectable population structure in both species, consistent with regional genetic connectivity across the sampled banks. In contrast, otolith chemistry revealed species-specific and bank-specific environmental variation, indicating that individuals experienced localized habitat conditions not captured by genetic markers alone. This contrast shows that regional genetic connectivity can coexist with fine-scale ecological and demographic structuring.

Species-specific otolith patterns suggest that Red Snapper and Vermilion Snapper use or experience shelf-edge habitats differently within this connected system. Red Snapper showed otolith patterns consistent with more localized reef-associated habitat use, whereas Vermilion Snapper exhibited a broader elemental fingerprint consistent with exposure to more heterogeneous environmental conditions. Together with differences in age structure and growth, these results indicate that genetically connected banks may still differ in habitat use, demographic composition, and local ecological conditions relevant to resilience.

From a management perspective, these findings are consistent with current regional stock treatment in the northern Gulf and also show that genetically connected stocks can contain ecologically meaningful spatial heterogeneity. As ocean conditions continue to change, integrating genomics with otolith-based approaches provides a framework for detecting shifts in habitat quality, recruitment pathways, and connectivity across shallow and mesophotic shelf-edge ecosystems. This multi-marker perspective can help strengthen adaptive management by distinguishing broad genetic connectivity from finer-scale ecological processes that may influence the resilience of snapper fisheries.

## Supporting information

Supplementary_Figures_Snappers_Habitat_Connectivity_Roa-Varon_etal_2026

Supplementary_Otolith_Materials-and_Methods_Snappers_Habitat_Connectivity_Roa-Varon_etal_2026

Supplementary_Tables_Snappers_Habitat_Connectivity_Roa-Varon_etal_2026

## Acknowledgements

The authors thank the captain and the NOAA R/V *Southern Journey* crew, Matt Galaska, Will Jenkins, Jonathan Quigley, Jen Herting, and Jayci Grosso for their support in collecting the samples. Thanks to Siyah Yongue, Meredith Anderson, Caroline Caron, Sheila Faith Linville, and Clara Demopoulos for assistance in the laboratory preparation of otoliths. Jane Rudebusch created the locations map (USGS). Gretchen Swarr trained ARV on using the LA-ICPMS at Woods Hole Oceanographic Institution. Olivia Cheriton tailored the MATLAB script to our otolith dataset (USGS). Funding for this work was provided by NOAA’s National Centers for Coastal Ocean Science, Competitive Research Program, and Office of Ocean Exploration and Research under award NA18NOS4780166 to Lehigh University, and by a Lehigh University CAS Dean’s Opportunity Grant. Additional support was provided by **MOA-2019-077/11806 to the U.S. Geological Survey Wetland and Aquatic Research Center.** Finally, portions of the laboratory work and data analysis were conducted with support from the Laboratories of Analytical Biology (https://ror.org/05b8c0r92) at the National Museum of Natural History. Any use of trade, firm, or product names is for descriptive purposes only and does not imply endorsement of the U.S. Government.

## Author contributions

ARV performed genomic and otolith laser ablation lab work, curated the data, performed analyses, generated visualizations, wrote the original draft, and incorporated edits.

NP collected specimens, reviewed, and edited the manuscript.

AD collected specimens, reviewed, and edited the manuscript.

PMc performed otolith preparation and reviewed the manuscript.

LT performed aging analyses and reviewed the manuscript.

SH collected specimens and reviewed the manuscript.

AWD collected specimens and reviewed the manuscript.

SH secured funding for the project, conceptualized it, and reviewed the manuscript.

AMQ secured funding for the project, conceptualized it, reviewed, and edited the manuscript.

All authors contributed to the article and approved the submitted version.

## Ethics approval

This project was approved by the National Museum of Natural History’s (NMNH) Institutional Animal Care and Use Committee (IACUC; SI-21020), and all parts of this study have been conducted in accordance with the approved protocols and relevant guidelines and regulations.

## Data Availability

Raw data and supplementary files supporting this study have been deposited in Figshare as separate records.

## Supplementary Figures

private Link: https://figshare.com/s/5331a4ecc0e5d99db72b DOI: 10.25573/data.32826596

## Supplementary Otolith Materials & Methods

private Link: https://figshare.com/s/a9b3385669c2c8cf3196 DOI: 10.25573/data.32826617

## Supplementary Tables

private Link: https://figshare.com/s/518ee567c90cc2039efc DOI: 10.25573/data.32826608

## Otolith Microchemistry Raw Data

private Link: https://figshare.com/s/7d2cee21d1a1b04c6d34 DOI: 10.25573/data.29736206

