## Supplementary_Figures_Snappers_Habitat_Connectivity_Roa-Varon_etal_2026 for "High Connectivity, Local Signatures: Genomic and Otolith Evidence from Reef-Associated Snappers"

**Supplementary Figures**

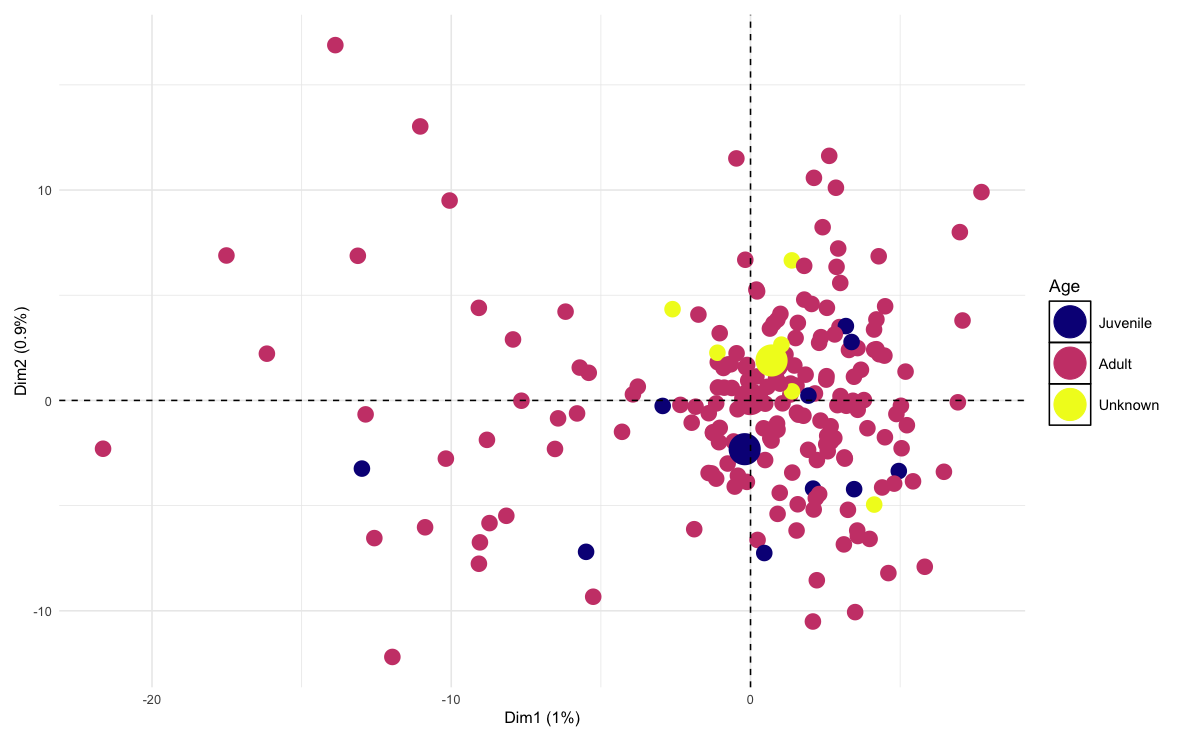
**A)**

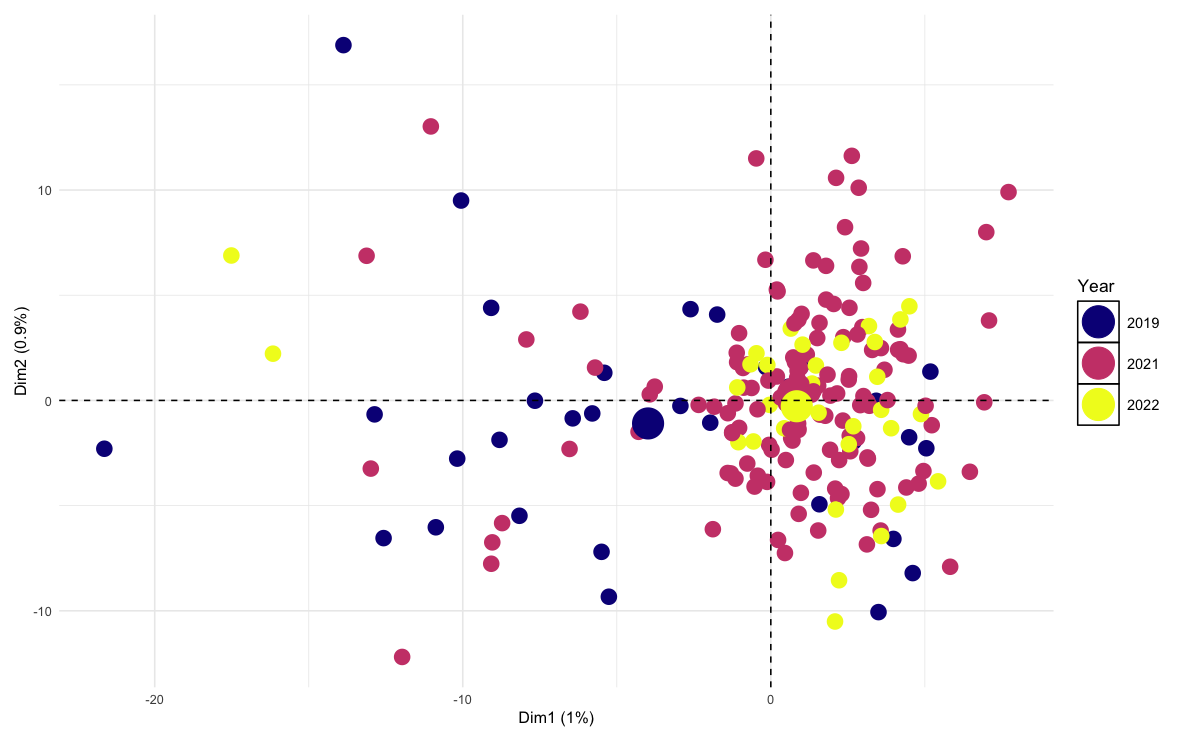
**B)**

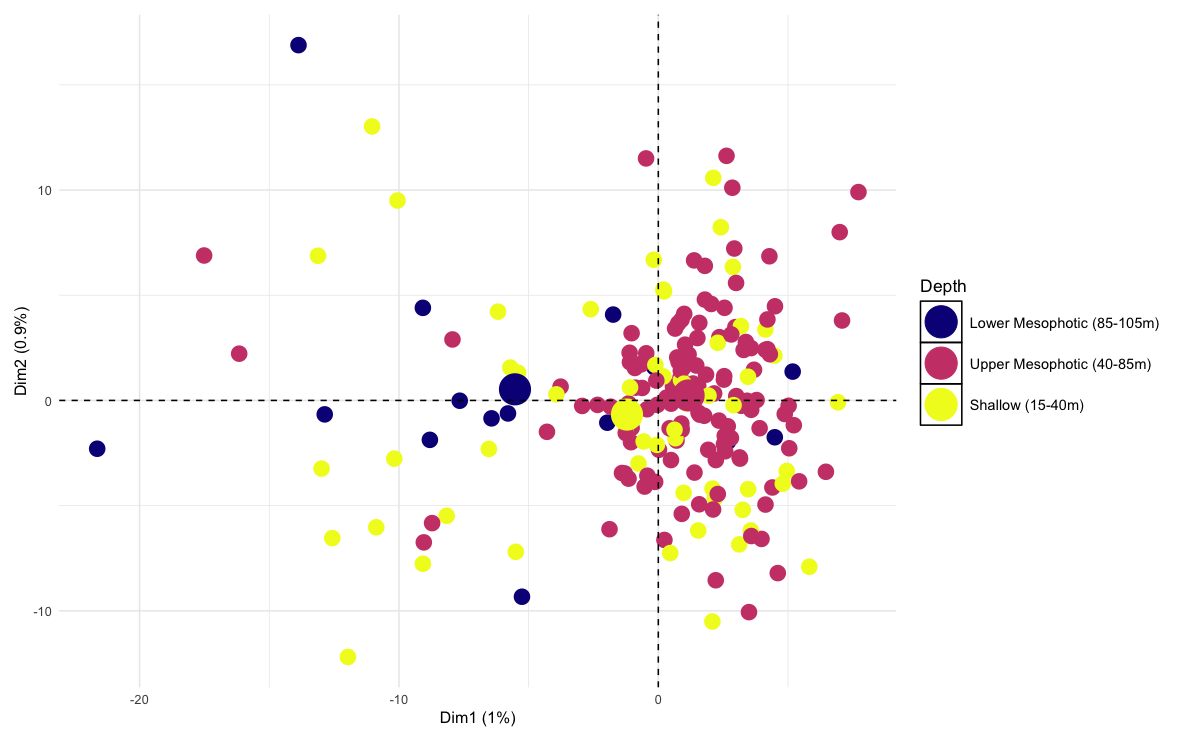
**C)**

**Figure S1.** Additional PCA projections for Red Snapper (*Lutjanus campechanus*) stratified by (A) age class, (B) sampling year, and (C) depth stratum. Individuals showed broad overlap among categories, with no evidence of discrete clustering.

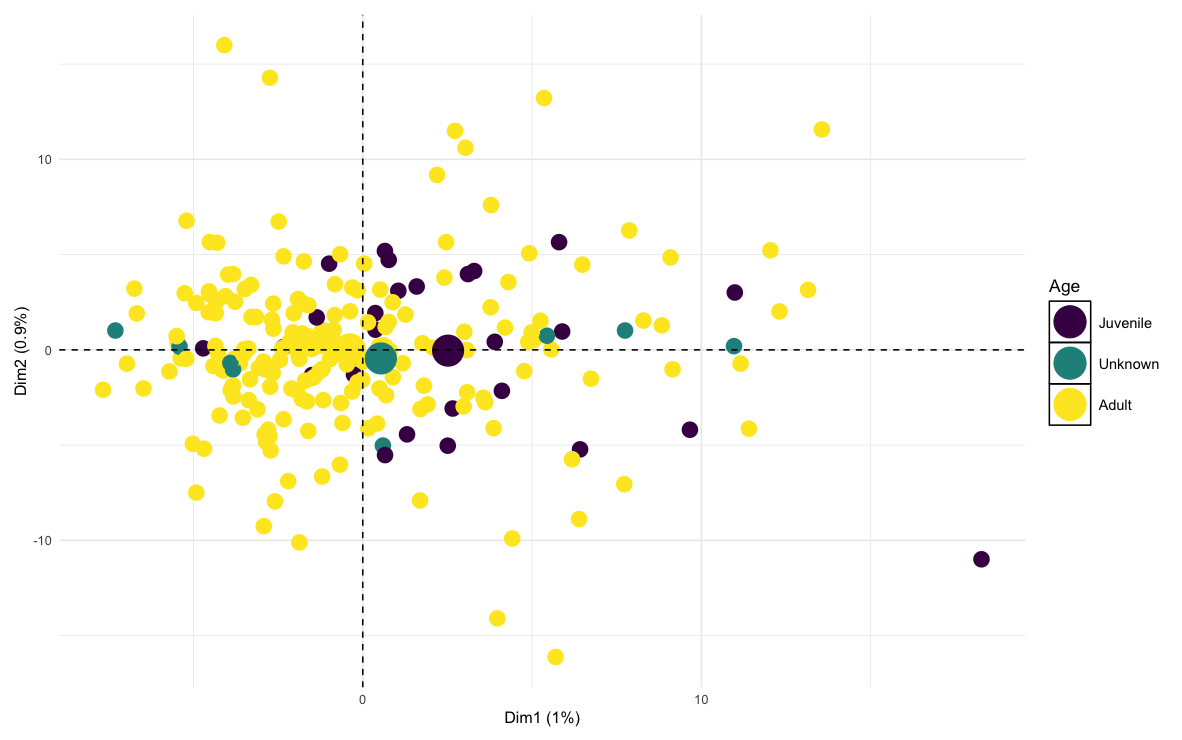
**A)**

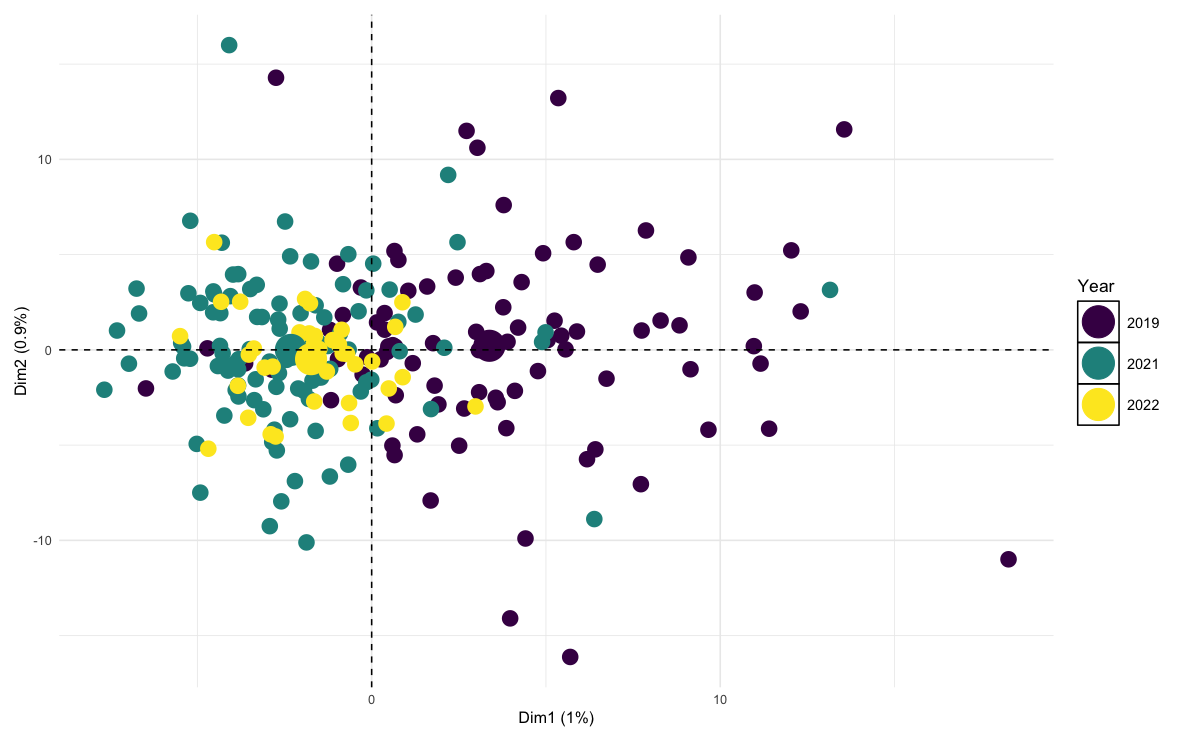
**B)**

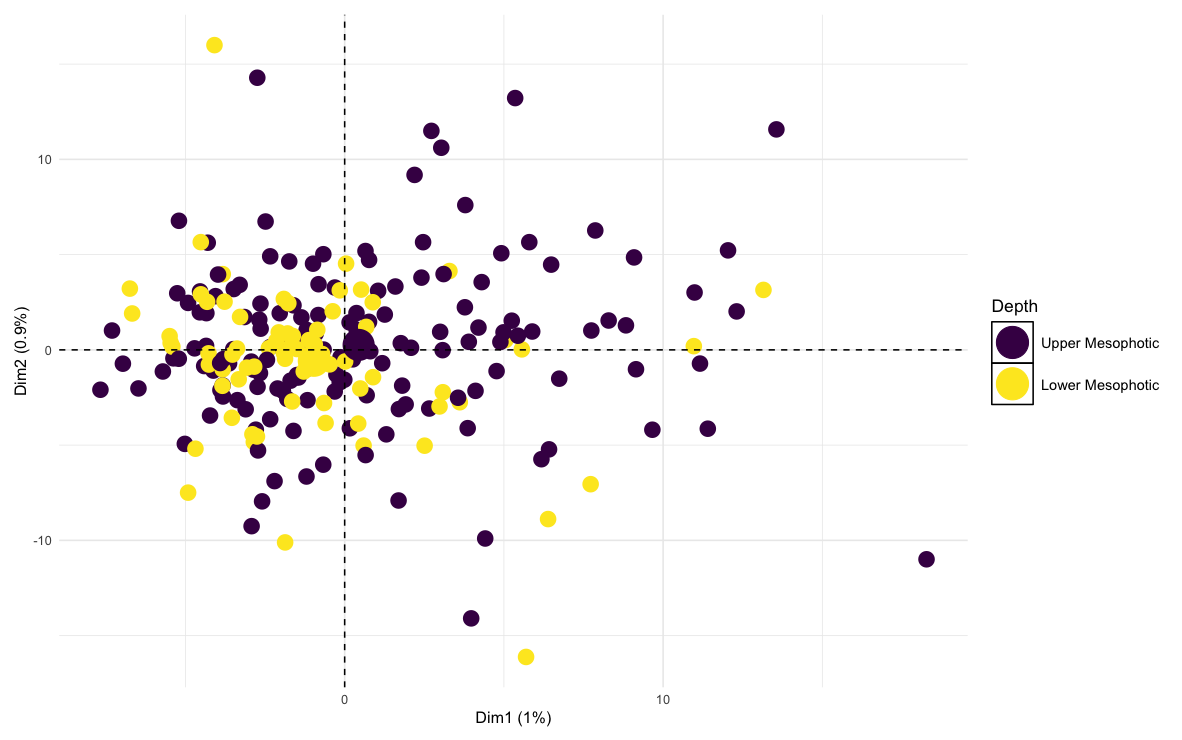
**C)**

**Figure S2.** Additional PCA projections for Vermilion Snapper (*Rhomboplites aurorubens*) stratified by (A) age class, B) sampling year, and (C) depth stratum. Individuals showed broad overlap among categories, with no evidence of discrete clustering.

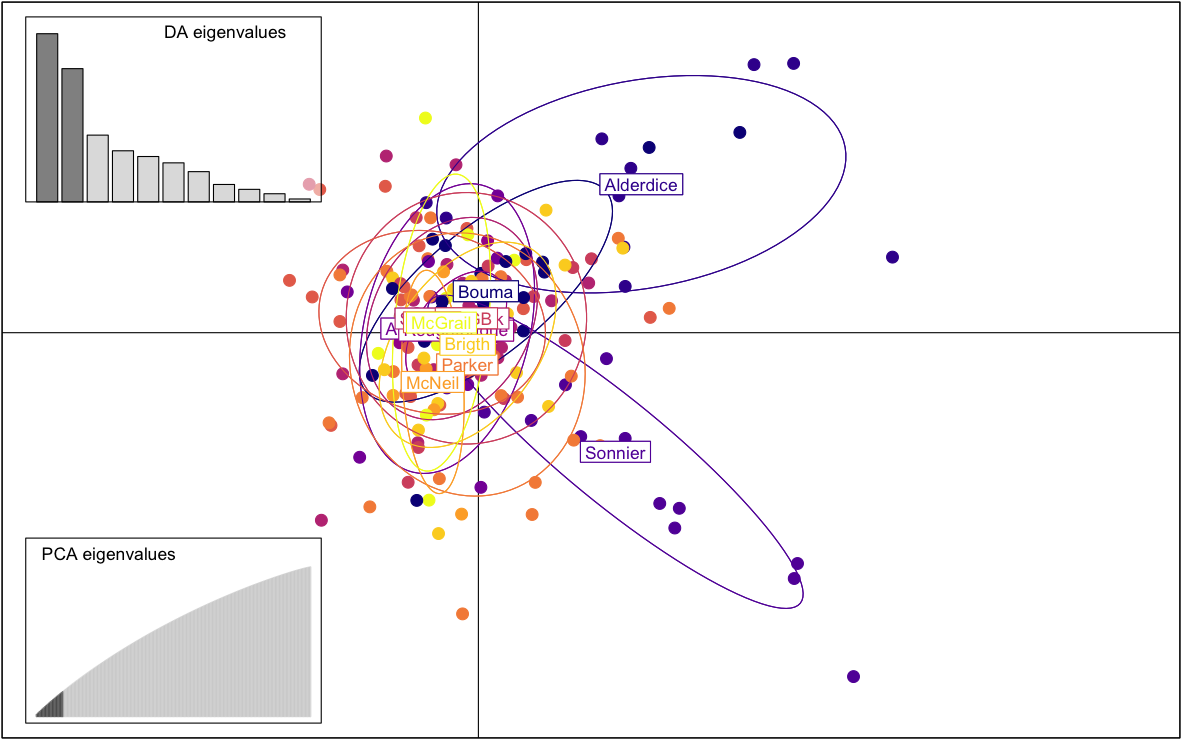
**A)**

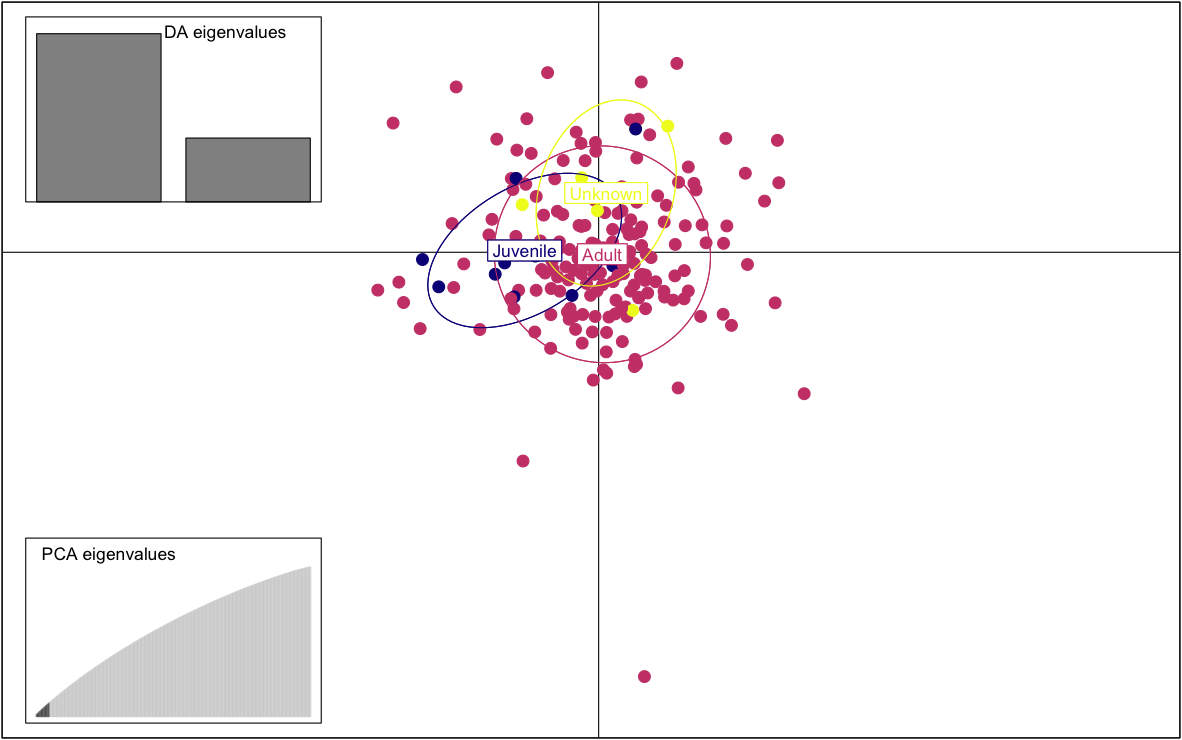

**B)**

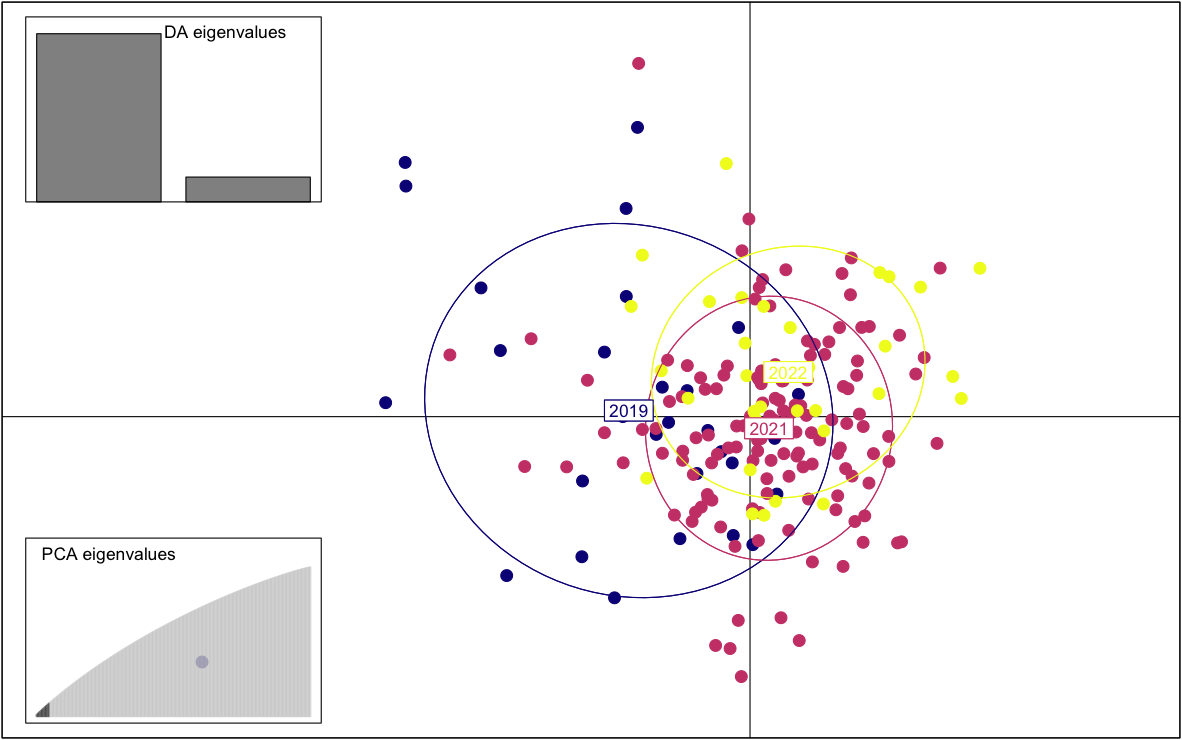
**C)**

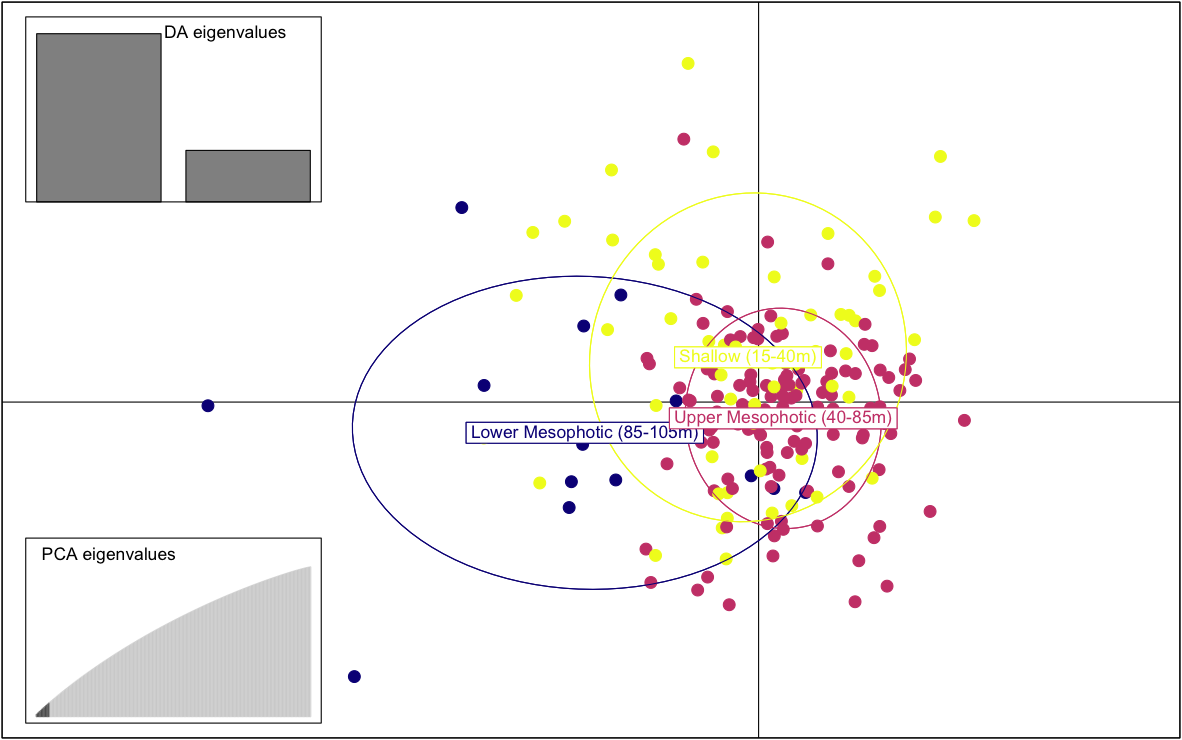

**D)**

**Figure S3.** Discriminant analyses of principal components (DAPC) for Red Snapper (*Lutjanus campechanus*) showing genetic structure across alternative grouping factors. Panels illustrate DAPC scatterplots with group ellipses, discriminant eigenvalue barplots, and cumulative variance explained for (A) sites, B) age classes, (C) sampling years, and (D) depth strata. Across all factors, individuals show extensive overlap in discriminant space, broadly distributed assignment probabilities, and a steep decline in discriminant eigenvalues followed by a long tail, indicating limited discriminatory power and weak genetic structure.

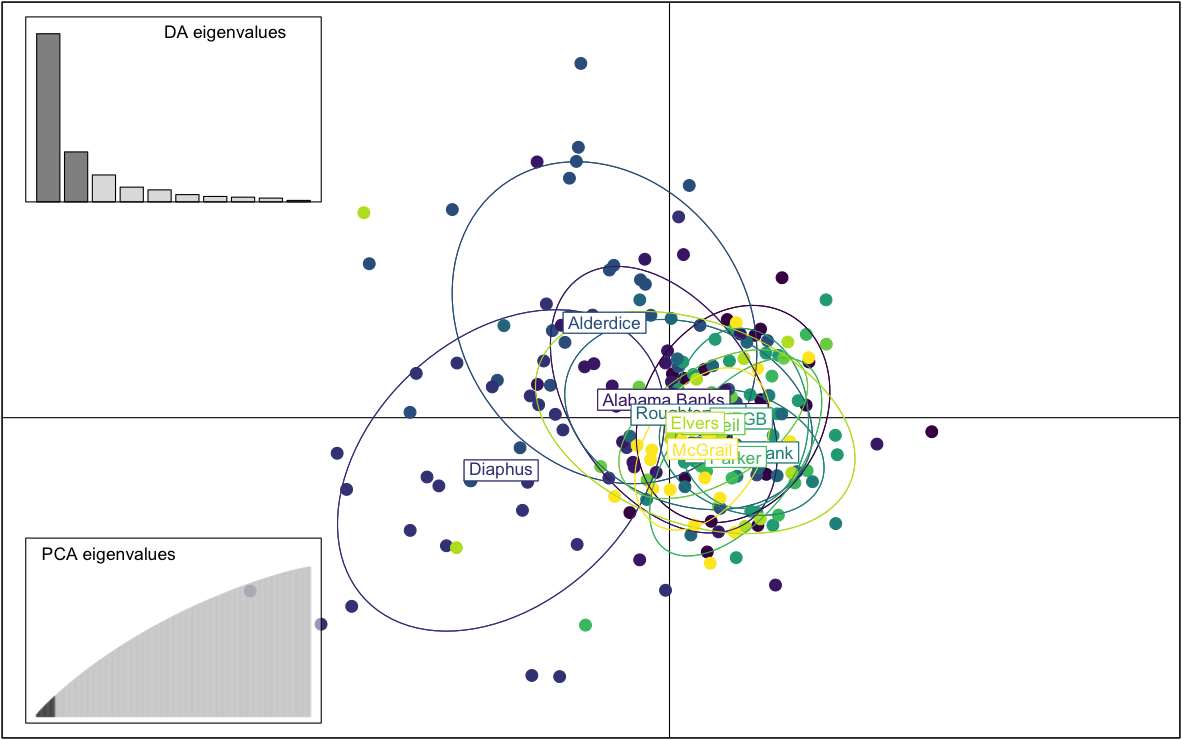
**A)**

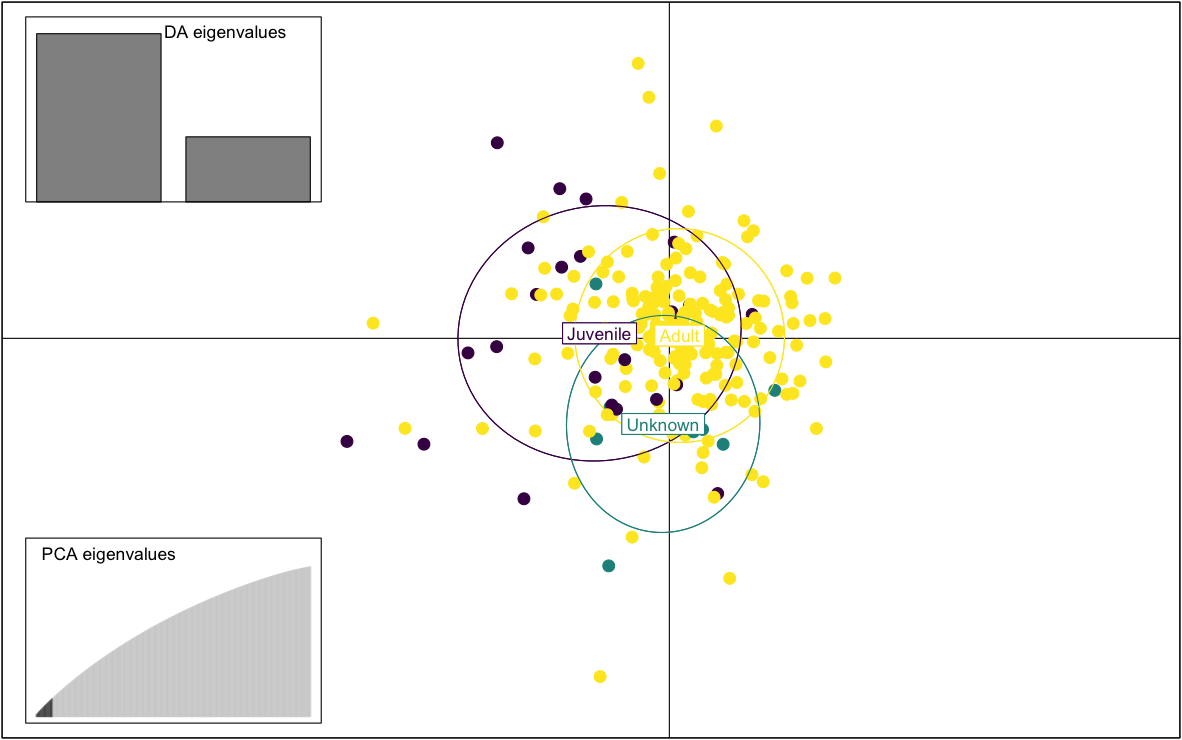

**B)**

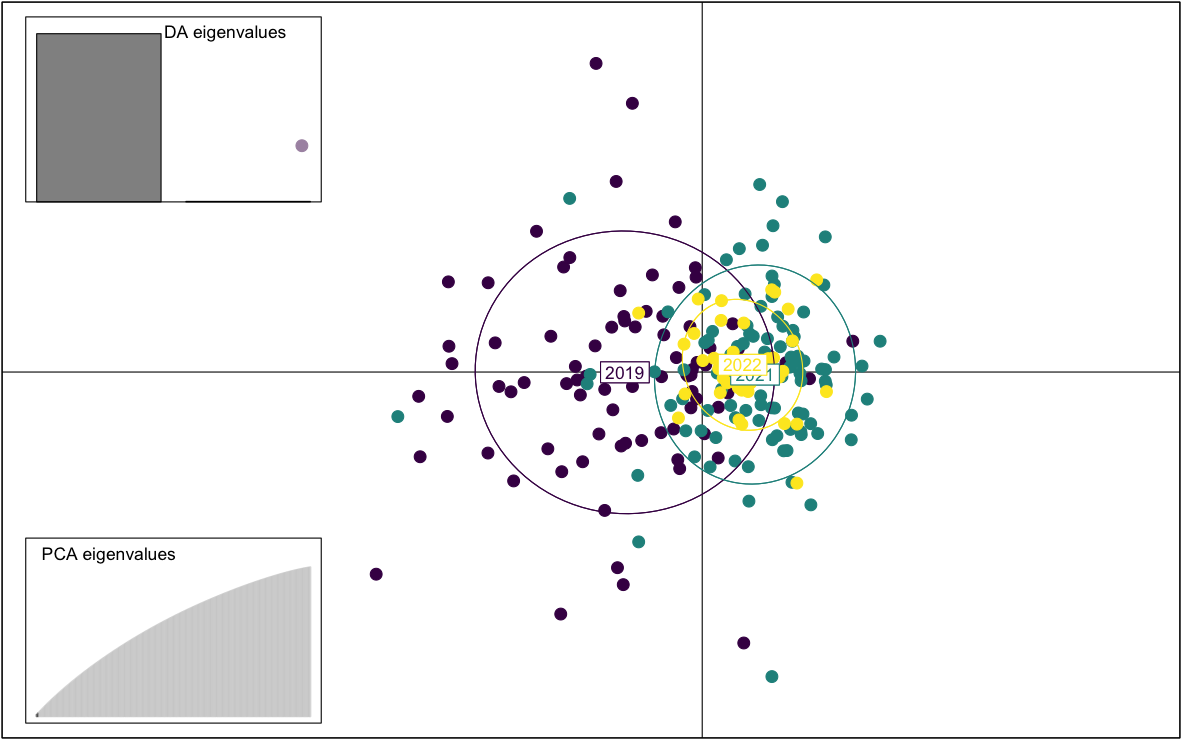
**C)**

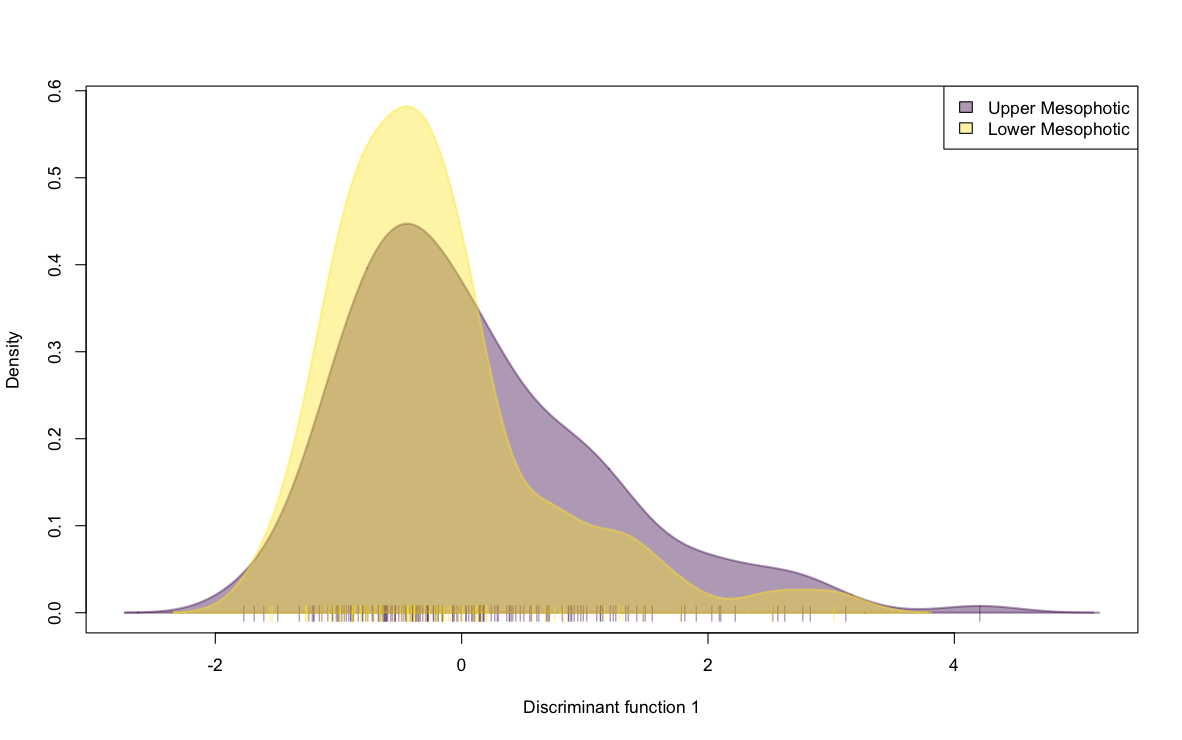
**D)**

**Figure S4.** Discriminant analyses of principal components (DAPC) for Vermilion Snapper (*Rhomboplites aurorubens*) showing genetic structure across alternative grouping factors. Panels A–C present DAPC scatterplots with group ellipses, discriminant eigenvalue barplots, and cumulative variance explained for individuals grouped by (A) site, (B) age class, and (C) sampling year. (D) Because depth included only two strata, only one discriminant function was retained; therefore, the distributions of individual scores along that function are shown for the upper- and lower-mesophotic strata.

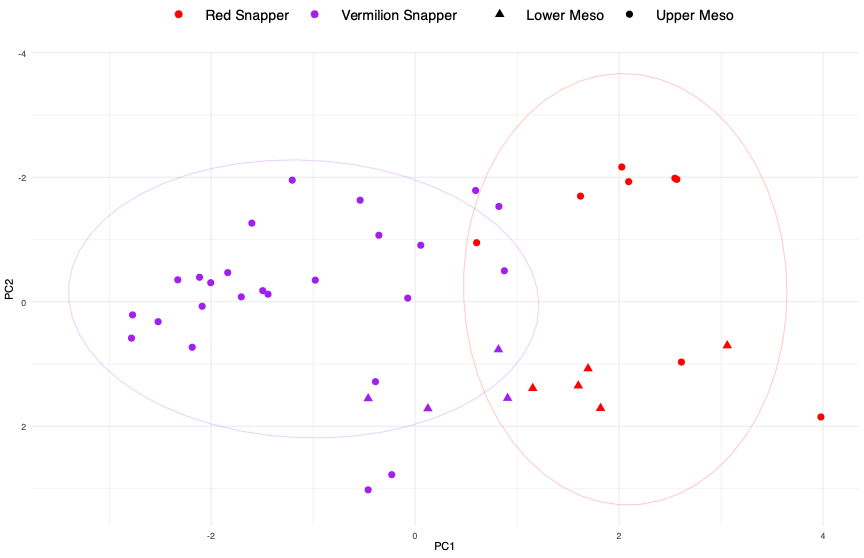

**Figure S5.** Exploratory PCA of otolith elemental composition for Red Snapper (*Lutjanus campechanus*) and Vermilion Snapper (*Rhomboplites aurorubens*) at West Flowers Garden Bank (WFGB), the only site with both species represented across Upper Mesophotic and Lower Mesophotic. Points are coded by species and depth stratum, and ellipses summarize group dispersion.

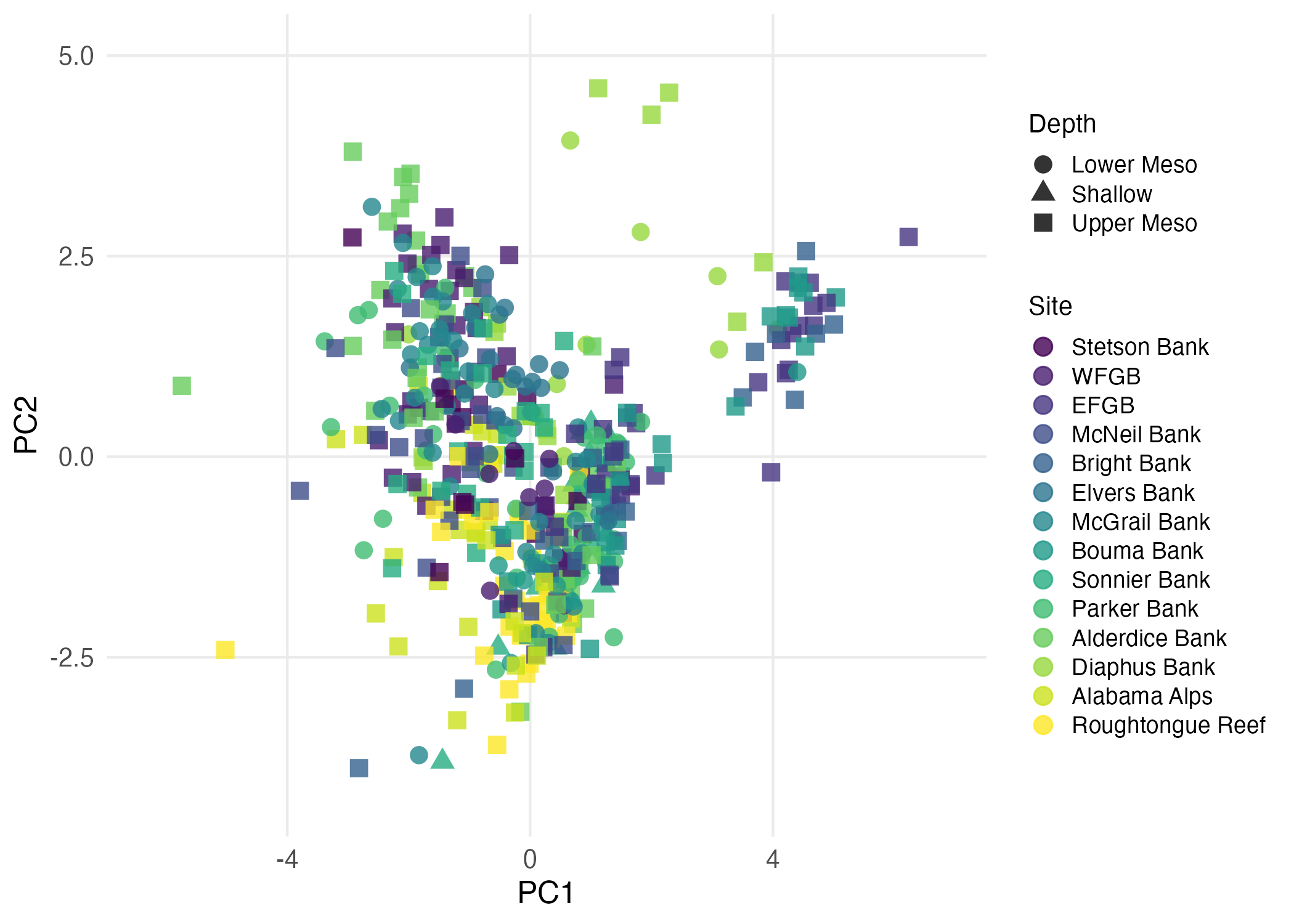

**Figure S6.** Principal component analysis of scaled otolith elemental composition colored by depth strata. Shallow, upper mesophotic, and lower mesophotic samples show extensive overlap in multielement space, indicating weak depth-associated structure in otolith chemistry.
