## Supplementary_Otolith_Materials-and_Methods_Snappers_Habitat_Connectivity_Roa-Varon_etal_2026 for "High Connectivity, Local Signatures: Genomic and Otolith Evidence from Reef-Associated Snappers"

**Supplementary Otolith Materials and Methods**

**Otoliths Preparation** Red Snapper (*Lutjanus campechanus*) and Vermilion Snapper (*Rhomboplites aurorubens*) were collected from shelf-edge banks within the Flower Garden Banks National Marine Sanctuary (FGBNMS) and neighboring sites in the northwestern Gulf of America across shallow (15-40 m), upper mesophotic (40-85 m), and lower mesophotic (85-150 m) depth zones using standardized hook-and-line fishing methods. Collections were conducted in 2019, 2021, and 2022 aboard the R/V *Southern Journey*. Sagittal otoliths were extracted from 250 Red Snapper and 234 Vermilion Snapper ). Right-side otoliths were cleaned to remove any contaminants by scrubbing with a soft-bristled brush, sonicated for 5 min, triple-rinsed with ultrapure water, and allowed to dry in a laminar flow hood. Following the decontamination procedure, otoliths were handled only with acid-washed equipment, embedded in Epoxies, Etc™ (Cranston, Rhode Island, USA) 20-3068 epoxy resin, and allowed to set for a minimum of three days. A thin (0.5 mm) and a thicker (1.5 mm) transverse section were cut through the primordium using a Buehler (Lake Bluff, IL, USA) IsoMet® low speed saw and mounted on glass slides using CrystalBond™ (Aremco Products, Inc., Valley Cottage, NY, USA) 509 thermoplastic glue. Thin sections were processed for ageing (refer to Otolith Age Readings below), while thicker sections were polished, sonicated for 5 min, triple-rinsed with ultrapure water, and allowed to dry. Thicker sections were then removed from the glass slides, and thermoplastic glue was removed using acetone. Excess epoxy around the otolith section was carefully trimmed off, then otoliths were again sonicated for 5 min, triple-rinsed, and allowed to dry. Sections were carefully placed core-side-down onto Buehler® MetGrip Liner adhesive within Buehler® Plastic Ring Forms (1 inch). After all otoliths were placed, the plastic rings were filled with epoxy resin to create ‘pucks’, allowed to set for three days, then removed from the adhesive liner and polished to ensure all otolith sections were exposed on an even plane. The pucks were then analyzed via Laser Ablation–Inductively Coupled Plasma–Mass Spectrometry (LA-ICP-MS).

**Otoliths Laser Ablation** The LA-ICPMS operating parameters for the laser were as follows: 90% power, 10 Hz pulse rate, and an ablation spot size of 64 μm, translated across the surface of the otoliths at a scan speed of 10 μm s ¹. A 20-second washout was performed after each sample to collect background counts.

We collected data on 12 analytes: lithium (7Li), boron (11B), magnesium (24Mg), calcium (43Ca and 48Ca), manganese (55Mn), cobalt (59Co), copper (63Cu), strontium (86Sr and 88Sr), and barium (137Ba and 138Ba). NIST 612 served as the standard, while MACS-3 was monitored as an internal standard. Both standards were processed at the beginning and end of each sample run (approximately 10-12 otoliths). A time-varying standard was established using simple linear regression between the mean of the first NIST 612 standard region and the mean of the NIST 612 standard region at the end of the sample run. Additionally, background values were determined for each element by using the output signal prior to running the first standards and after the last standard. Similar to the standards, a time-varying background was identified through linear regression between the mean of the first background region and that of the second. The time-varying background was then subtracted from each element's concentrations. Subsequently, the resulting trace element concentrations were analyzed as ratios with 43Ca, and the data were converted to concentration ratios based on measurements of the NIST 612 standard using the following formulation:

$$\frac{mmol}{molCa}=\left( \frac{1}{k} \right)*\left( \frac{element-background}{43Ca-background} \right)$$

Where k is a correction factor obtained by dividing the time-varying standard (measured NIST 612 glass ratios) by the mean element:Ca ratios (ppm) reported in Jochum et al. (2011). The resultant elemental ratios are presented in units of mmol/mol.

**Otolith Age Readings** Transverse cuts prepared on slides were sent to the National Marine Fisheries Service, Panama City Laboratory. To confidently assign annuli counts to red snapper and vermilion snapper, the ideal thickness of the transverse cut is approximately 0.5 mm. The cuts were thinned using 600 grit sandpaper on a Crystalite, Inc. (Everett, WA, USA)CrystalMaster™ lapidary grinder. Buehler MicroPolish® II alumina 1µm was used after sanding to polish the otolith surface with a Buehler MicroCloth polishing cloth applied to another lapidary grinder. Lastly, a thin coat of CytoSeal™ (Epredia, Portsmouth, NH, USA) was applied to the thinned slide to protect the otolith and enhance the appearance of the section. Age readings were estimated from annuli (opaque zone) counts and the degree of marginal edge completion, viewed under transmitted and/or reflected light. Typically, marine fish inhabiting the waters around the southeastern U.S. complete annulus formation (opaque zone formation) by late spring to early summer (Johnson, 1983; Patterson et al., 2001; Wilson and Nieland, 2001; Garcia et al., 2003; White and Palmer, 2004; Allman et al., 2005). For instance, an otolith with two completed annuli and a large translucent zone (≥66% of the preceding translucent zone width) would be classified as age 3 if the catch date coincides with January 1st through June 30th, as the third annulus would be presumed to have soon formed. After June 30th, the calendar age equaled the annuli count. By this traditional method, an annual age cohort is based on a calendar year rather than time since spawning (Jearld, 1983; Vanderkooy et al., 2020). Vermilion snapper otolith sections tend to have a varied appearance, making it difficult to confidently count opaque zones (Allman et al., 2001).

**Collinearity and Variance Inflation Factor** Because individual otolith elements may covary due to the shared environmental drivers (e.g., water mass, chemistry, salinity) and/or physiological processes (e.g., redox metabolism, trace metal binding, growth), we quantified collinearity to evaluate the redundancy and to identify multielement “families” within and across species. For each species, Pearson correlation matrices were calculated using pairwise complete observations, and hierarchical clustering (correlation distance, complete linkage) was used to visualize elemental modules (Dormann et al., 2013, James et al., 2013). To further quantify multivariate redundancy, variance inflation factors (VIFs) were calculated using a regression-based approach in which each element was modeled as a function of the remaining elements and VIFs computed as 1/(1 - R^2^) (James et al. 2013). We interpreted VIF <5 as low-moderate collinearity, 5-10 as moderate-high, and >10 as strong collinearity. This framework allowed us (i) to assess the dimensional independence of geochemical and physiological axes, (ii) determine whether multivariate analyses (PCA, CAP) were justified without variable reduction, and (iii) to compare species-specific differences in otolith elemental architecture.
